# Resource: A Compendium of HLA-Type and Expression in Pediatric Cancer Models

**DOI:** 10.1101/2025.07.24.666682

**Authors:** Yiwen Guan, Ishika Mahajan, Vikesh Ajith, Isaac Woodhouse, Tima Shamekhi, Pouya Faridi, Ron Firestein, Claire Xin Sun

**Affiliations:** Centre for Cancer Research, Hudson Institute of Medical Research, Clayton, Australia. Department of Medicine, School of Clinical Sciences, Monash University, Clayton, Australia; Department of Medicine, School of Clinical Sciences, Monash University, Clayton, Australia; Monash Proteomics and Metabolomics Platform, School of Clinical Sciences, Monash University, Clayton, Australia

**Keywords:** HLA, Immunogenicity, pediatric cancer, data recourse

## Abstract

Cancer immunotherapy has transformed cancer treatment by harnessing the immune system’s natural ability to recognize and eliminate tumor cells, offering an effective and less toxic alternative particularly suitable for pediatric patients. Advances such as immune checkpoint inhibitors targeting PD-1 and CTLA-4, successful adoptive cell therapies like CAR T-cell treatments, and innovative cancer vaccines have notably improved outcomes across various cancer types. Central to immunotherapy is antigen presentation, where tumor-derived peptides loaded onto the Major Histocompatibility Complex molecules facilitate immune recognition. Precise HLA typing is crucial, especially in pediatric cancer research, where cell lines serve as essential preclinical models. The Childhood Cancer Model Atlas (CCMA) initiative provides comprehensive profiling, including HLA typing and neoantigen predictions for 287 cancer cell lines. This publicly accessible resource enables researchers to better understand immune-cancer interactions, advance personalized immunotherapies, and develop targeted, less toxic treatments specifically beneficial for children with cancer.

## Introduction

Cancer immunotherapy represents a transformative approach to cancer treatment, harnessing and enhancing the body’s immune system to recognize and eliminate tumor cells. Recent advancements, including immune checkpoint inhibitors targeting PD-1 and CTLA-4 in melanoma and lung cancer, the remarkable success of adoptive cell therapies such as CAR T-cell therapy in blood cancers, and developments in cancer vaccines, have significantly improved patient outcomes across various cancer types^1,2^. Moreover, advances in cancer vaccines have elicited *de novo* T-cell responses against tumor antigens, offering further promise in combating diverse malignancies^3^.

Antigen presentation plays a central role in immune recognition, acting as a bridge between tumor cells and the immune system. This process begins with intracellular proteins being degraded into peptides by the proteasome, followed by the loading of these peptides onto major histocompatibility complex (MHC, known as HLA in human) molecules encoded by the Human Leukocyte Antigen (HLA) genes. HLA typing is a critical aspect of immunogenetics, especially in the context of cancer research and immunotherapy development. HLA molecules, encoded by genes on chromosome 6, are essential in the presentation of antigens to T-cells, thus influencing immune recognition and tumor evasion. In pediatric cancer, where cell lines serve as valuable models for studying tumor biology and therapeutic response, accurate HLA typing is a critical part for researchers to investigate the interaction between cancer cells and the immune system.

Cell lines serve as essential preclinical models and powerful tools in the development of innovative immunotherapy treatments. The Childhood Cancer Model Atlas (CCMA) initiative is at the forefront of pediatric cancer research, providing extensive multi-omics profiling of 287 cancer cell line models. The integration of comprehensive HLA typing predicted neoepitope profiles derived from missense, in-frame indels, frameshift mutations, and assessments of immunogenic activity provides valuable insights into the immunological characteristics of these cell lines, thereby advancing personalized medicine strategies. Researchers can leverage these data to better anticipate immune responses against cancer neoantigens and evaluate the potential effectiveness of immunotherapies, including checkpoint inhibitors, T-cell therapies, and cancer vaccines. Additionally, the availability of extensively HLA-typed cell lines greatly facilitates collaborative, large-scale immunogenomic studies, enabling deeper exploration into the influence of HLA diversity on therapeutic responses, and mechanisms underlying treatment resistance.

This extensive HLA characterization of pediatric cancer models, publicly accessible via the CCMA dashboard (https://vicpcc.org.au/dashboard), will enhance the ability to test immune-based therapies and improve our understanding of immune-cancer interactions, ultimately contributing to the development of novel, targeted treatments for childhood cancers.

## Results

### The annotation of HLA types for the CCMA

The CCMA hosts the largest single-site collection of pediatric cancer cell line models. The 287 cell lines reported in this study span nine cancer types, including pediatric high-grade gliomas (pHGG, n=114), which consist of 67 diffuse midline glioma with H3K27M (DMG-H3K27M), 35 high-grade glioma with wildtype H3 (HGG-H3WT), 10 diffuse hemispheric glioma with H3G34R/V mutation (DHG-H3G34), 1 astrocytoma and 1 pineoblastoma. Bone and soft tissue sarcomas (n=33) include 12 ewing’s sarcoma, 11 osteosarcoma, 7 rhabdomyosarcoma, 2 chordoma, 1 inflammatory myofibroblastic tumor (IMFT). Embryonal tumors (n=35) include 20 atypical teratoid/rhabdoid tumor (ATRT), 11 medulloblastoma, 2 embryonal tumor with multilayered rosettes (ETMR), 2 malignant rhabdoid tumor (MRT). Ependymal tumors (n=9) include 9 ependymoma and neuroendocrine tumors (n=17) include 17 neuroblastoma. Other rare cancer types (n=4) consist of 2 central nervous system sarcoma, 1 circumscribed astrocytic glioma and 1 nerve sheath tumor. Additionally, there are 59 tumor-associated non-malignant cell lines, and 16 adult high-grade gliomas (aHGG) included in the cohort. Among the 287 unique cell lines, 174 have been previously studied, while 113 are newly established models sourced from local hospitals and collaborators. New RNA-seq data (n=103) and WGS data (n=93) is presented in this study along with previously published CCMA datasets^4^.

Using high-resolution HLA typing and WGS/RNA-seq HLA typing with Optitype^5^, we determined 4-digit class I HLA types and characterized the level of HLA expression and antigen presentation pathways for 287 CCMA cell lines (Figure 1A; Table S1.1). Additionally, neoepitopes were predicted for each model using pVACseq^6^ based on the somatic variants identified by WGS (Figure 1A). The reported HLA types were determined using a robust method illustrated in Figure 1B. A total of 29 cell lines were assessed using the NGS high-resolution HLA typing assay, deemed as the gold standard. Of the remaining 258 lines, 206 had both WGS and RNA-seq data available for typing, 17 had only WGS-based inference, and 35 had only RNA-seq based inference. Amongst the 52 lines with single source typing results, the optimal solution was reported (Solution 0 from Optitype algorithm). Of the remaining 206 models with dual-source results, 154 showed complete concordance between the optimal WGS and RNA-seq typing solutions (Solution 0). In 24 models, homozygous HLA types were identified in RNA-seq, while the corresponding WGS types were heterozygous (more details in the next section). In these cases, the WGS Solution 0 results were reported. The remaining 28 models exhibited non-concurrent HLA typing, and their HLA types were subject to manual curation. Among these, 13 models achieved matching HLA types at 2-digit resolution (Table S1.2). The mean concordance rate of all the HLA alleles between the 28 mismatched cell lines was 82.14% at 2-digit resolutions and 69.05% at 4-digit resolutions (Figure S1A). Accurate determination of HLA types requires validation using high-resolution typing methods.

**Figure 1.**
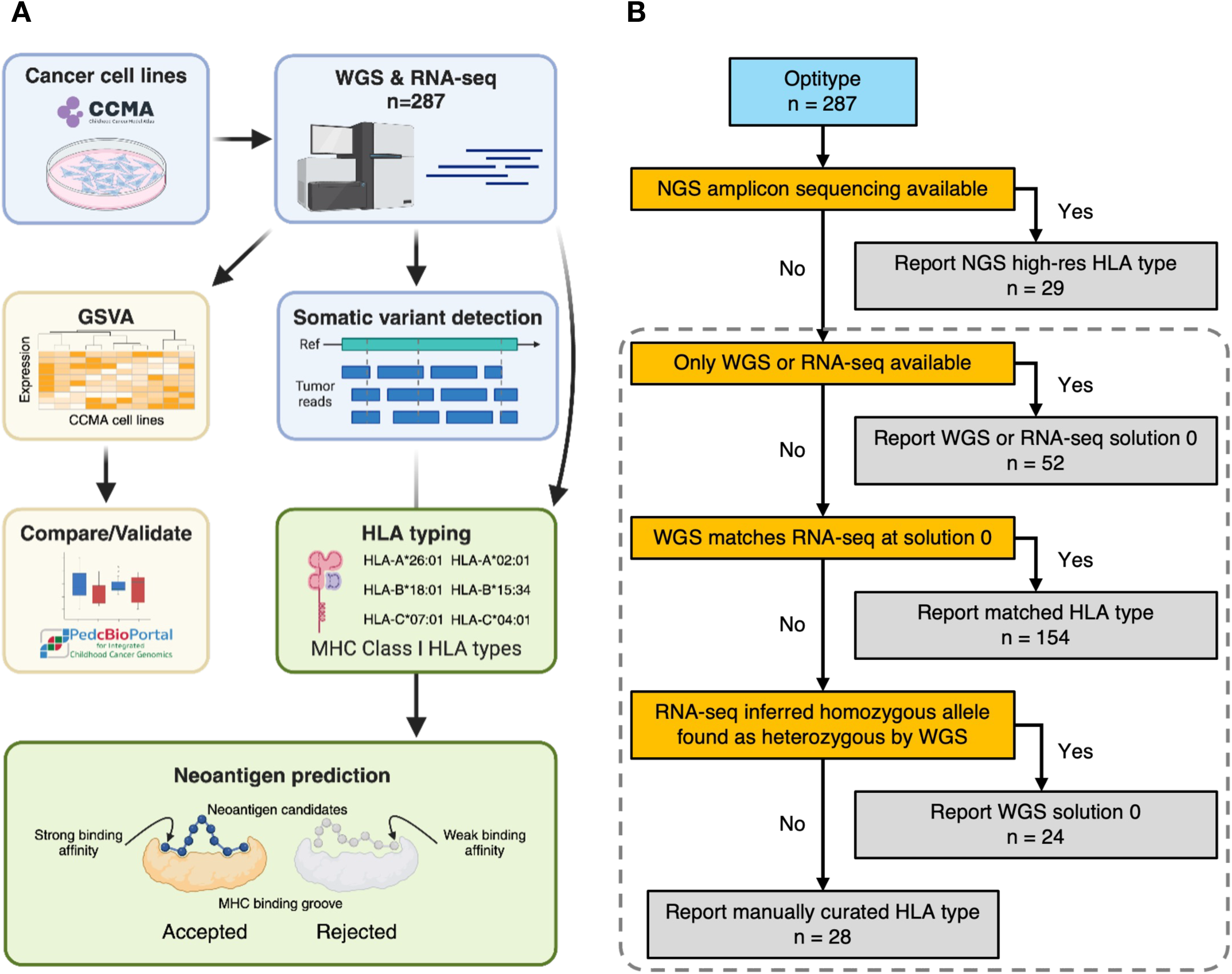
Computational workflow and data integration for HLA expression, typing, and neoepitope prediction in the CCMA. (A) WGS and RNA-Seq data from 287 cancer cell lines were processed for GSVA and Class I HLA typing. HLA pathway activity scores were compared to primary tumors (OpenPedCan). The cell line-specific somatic mutations and HLA types were processed to predict potentially effective neoepitopes. (B) Decision-making process for selecting the optimal HLA types. The final HLA types were determined hierarchically by each criterion based on data availability. HLA types inferred from Optitype were denoted by the dash-line box.

### The HLA supertypes and zygosity for the CCMA models

HLA supertypes represent distinct groups of HLA alleles that share peptide-binding specificities^7,8^. These supertypes have been widely used in cancer antigen identification^9–11^, neoantigen prediction^6^, and disease association studies^12,13^. The CCMA cohort exhibited a strong representation of HLA supertypes, which is important in immunotherapy studies due to their broad population coverage (Figure 2). Supertypes for HLA genes A and B were categorized for all cell lines. The most prevalent supertypes in the examined cell lines were A02, A03, and A01 for gene HLA-A (173, 166, and 132 of 572 alleles, respectively) and B07 and B44 for gene B (214 and 141 of 574 alleles, respectively) (Figure 2; Figure 3A).

**Figure 2.**
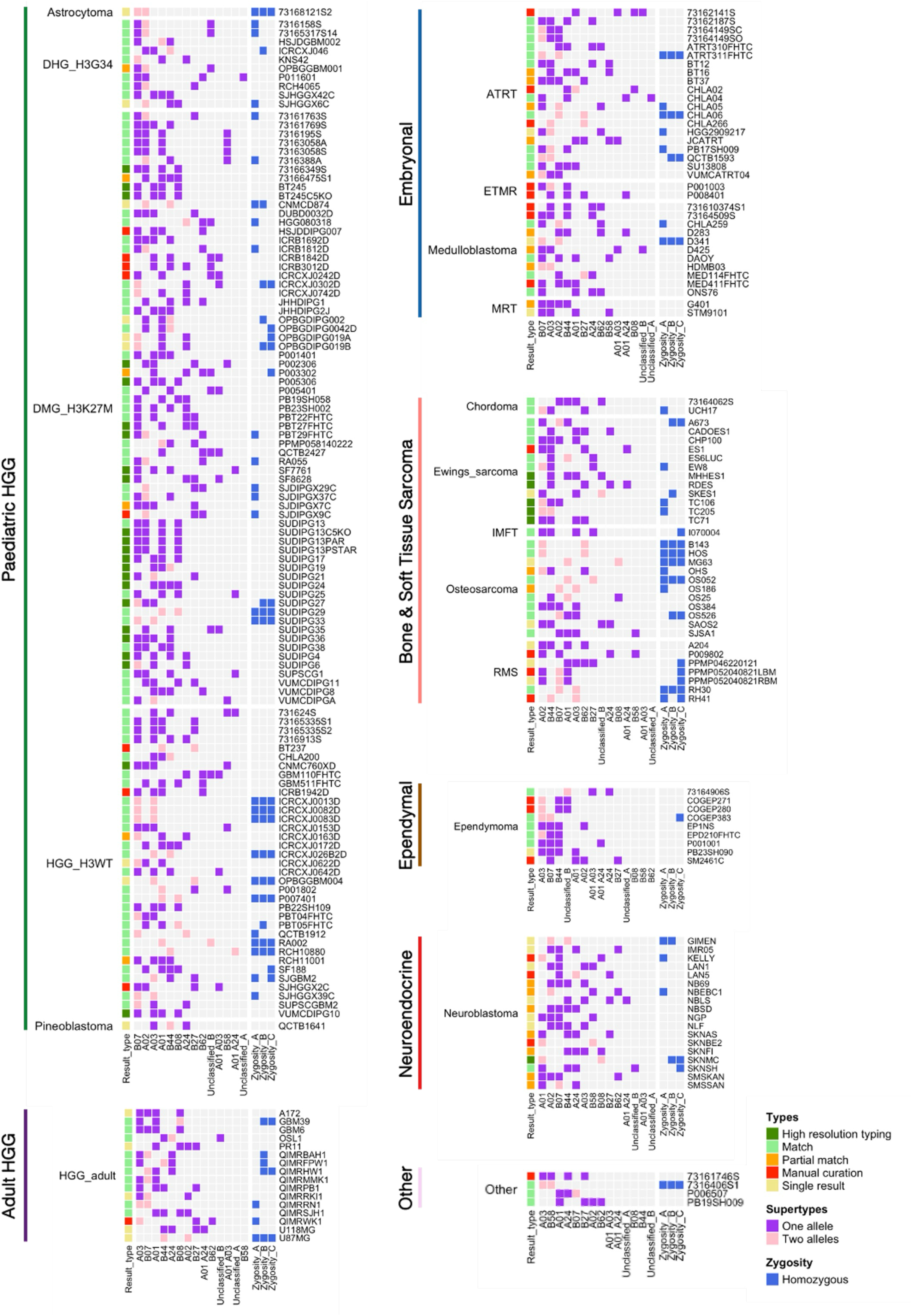
Tile plot for the CCMA HLA types. Tile plot showing data integration, supertypes, and zygosity for the CCMA models. Subplots were grouped by cancer type and clustered by CCMA cancer class. Data including result type (High-resolution typing, Match, Partial match, Manual curation, and Single result), supertypes (“One” representing allele and “Two” representing alleles), and zygosity (Homozygous) for each model were color-coded as indicated.

**Figure 3.**
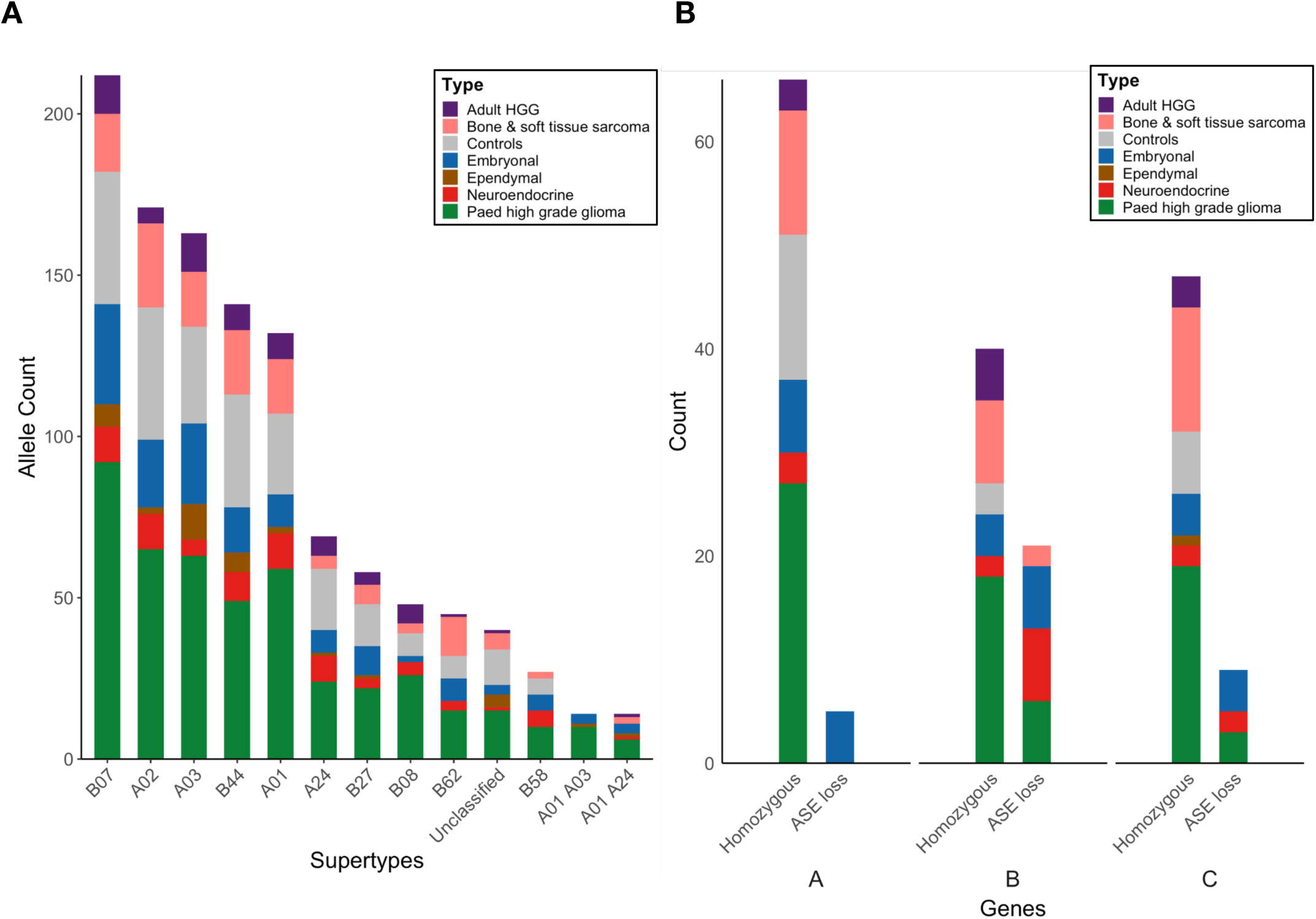
Distribution of HLA supertypes, zygosity, and ASE loss events across the CCMA. (A) Stacked bar chart showing the HLA allele count of CCMA models across 12 supertypes or unclassified. HLA supertypes were defined according to Sidney et al., 2008^8^. (B) The number of CCMA models with homozygous loci and ASE loss across Class I HLA genes. ASE loss events were inferred from comparing WGS and RNA-Seq HLA types. Twenty-one CCMA cancer classes were color-coded as indicated.

Heterozygous HLA sequences provide an advantage by enabling the presentation of diverse immunopeptides^14,15^. In contrast, homozygous HLA genotypes are associated with poorer clinical outcomes in cancer immune checkpoint blockade (ICB) therapy^16,17^. Recent studies revealed that HLA loss of heterozygosity (LOH) is a common cancer immune evasion strategy^18,19^. HLA LOH can occur through irreversible alterations to the HLA locus in chromosome 6, or reversible HLA allele-specific expression loss (ASE loss) at the transcriptomic level^20^. Both irreversible HLA LOH and ASE loss are associated with poor therapy outcomes^16,21^. In the CCMA cohort, we observed a significant proportion of homozygous HLA types (Figure 2), which we suspect may be attributed to HLA LOH. To this end, we analyzed the homozygosity frequency of class I HLA genes across different cancer types in CCMA (Figure 3B). The homozygosity frequencies for gene HLA-A ranged from 0% in ependymoma to 55% (6 out of 11) in osteosarcoma models. For gene HLA-B, frequencies ranged from 0% in ependymoma to 45% (5 out of 11) in aHGG models, while HLA-C frequencies ranged from 0% in DHG-H3G34 to 63% (5 out of 8) in rhabdomyosarcoma models. Osteosarcoma and ependymoma models exhibited the highest and lowest homozygosity frequencies among all CCMA major cancer types, respectively.

Furthermore, we identified discrepancies between HLA types inferred from WGS and RNA-seq in 24 models. Specifically, these models exhibited heterozygous HLA types in WGS but homozygous HLA types in RNA-seq, with the homozygous allele present within the heterozygous type. We confirmed the homozygous HLA types by RNA-seq using another widely used tool, arcasHLA^22^. All HLA types were consistent with the results obtained from Optitype. We hypothesize HLA allele specific expression was suppressed at the transcriptomic level, suggesting ASE loss. The 24 models with potential ASE loss were distributed across 8 of the 21 CCMA classes (Figure 2). ASE loss was most common in osteosarcoma (7 out of 11; 64%), ATRT (9 out of 20; 45%), and medulloblastoma (5 out of 11; 45%), but less frequent in DMG-H3K27M, DHG-H3G34, HGG-H3WT, neuroblastoma, and rhabdomyosarcoma. For all observed HLA-I ASE loss events, 5 out of 24 (21%) involved HLA-A, 21 out of 24 (88%) involved HLA-B, and 9 out of 24 (38%) involved HLA-C.

To test the robustness of HLA typing with WGS and RNA-seq, we compared Optitype results to NGS high-resolution typing. From 29 lines with high-resolution typing results available, there were two sets of isogenic cell line families (BT245 and BT245C5KO; SUDIPG13, SUDIPG13C5KO, and SUDIPG13PSTAR). Only the parental models were considered for accuracy calculations (BT245 and SUDIPG13). Of the 26 unique models, 25 achieved at least 2-digit accuracy, with mean accuracies of 96.81% and 94.19% at 2-digit and 4-digit resolutions, respectively (Table S2.1; Figure S1B). Only one model VUMCDIPG10 (3 true heterozygous loci) were incorrectly predicted by Optitype as homozygous by WGS at the HLA-A, B and C locus. Therefore, this line was reported as undetermined. The overall zygosity prediction accuracy was 96.15% (Table S2.2; Figure S1C). Most cell lines had complete matches while four had minor mismatches at the four-digit resolution. Overall, WGS and RNA-seq based HLA inference achieved high accuracy comparable to the standard high-resolution NGS typing method.

### The predicted neoepitope profiles for the CCMA

Neoantigens are peptides derived from somatic mutations generated by tumor cells. Effective neoantigens presented by the MHC can activate CD8^+^ and CD4^+^ T cells, triggering tumor-specific immune responses^23^. Neoantigens serve as key targets in cancer immunotherapies and personalized treatments. While extensive neoantigen research has been conducted in adult cancers such as lung cancer^24^, pancreatic cancer^25^, and melanoma^26^, there remains a significant knowledge gap in pediatric cancers. Using the pVACseq epitope pipeline with selected prediction algorithms and prioritization filters, we identified neoepitope candidates for 229 cell line models in the CCMA based on their WGS profiles. We employed a robust pipeline consisting of three binding prediction models (NetMHCpanBA, MHCflurry, and MHCnuggetsI), three elution prediction models (NetMHCpanEL, MHCflurryEL, and BigMHC-EL), and one immunogenicity prediction model (BigMHC-IM) to systematically prioritize neoepitopes. For initial exploratory analysis, we applied an IC50 binding threshold of <500 nM to filter candidates. The top neoepitope with the lowest IC50 score from any binding prediction model was selected for each variant to ensure the inclusion of all potentially effective neoepitopes.

Across the 229 models spanning seven cancer types, we identified a total of 12,811 putative neoepitope candidates with an IC50 score below 500 nM, while 7,028 (54.86%) were considered strong binders with an IC50 score below 50 nM (Figure 4A; Table S3.1). The distribution of neoepitopes per model was well-balanced across cancer types (Figure 4B). Models showed a median of 47 neoepitopes per sample. As expected, the non-malignant tumor associated cell line model showed the lowest number of predicted neoepitopes. To further incorporate the biological relevance of peptide processing, TAP-mediated peptide transport, and MHC presentation, we applied percentile filters (<2% and <0.5%) from elution prediction models to all neoepitopes meeting the IC50 <500 nM threshold. Out of 12,811 candidates, 7,315 (57.1%) and 4,417 (34.48%) neoepitopes had a percentile rank of <2% and <0.5%, respectively. Furthermore, we used peptide immunogenicity prediction models to assess whether these peptides were likely to trigger an immune response. Among all promising neoepitopes, 489 (3.82%) passed a BigMHC-IM score threshold of > 0.5. We observed 2,631 (20.54%) neoepitope candidates passing all binding and elution model filters, with 303 (2.37%) neoepitopes also passing the immunogenicity filter (Figure 4A). These stringent sets of peptides have higher likelihood of presentation, immunogenicity, and are prioritized for downstream testing.

**Figure 4.**
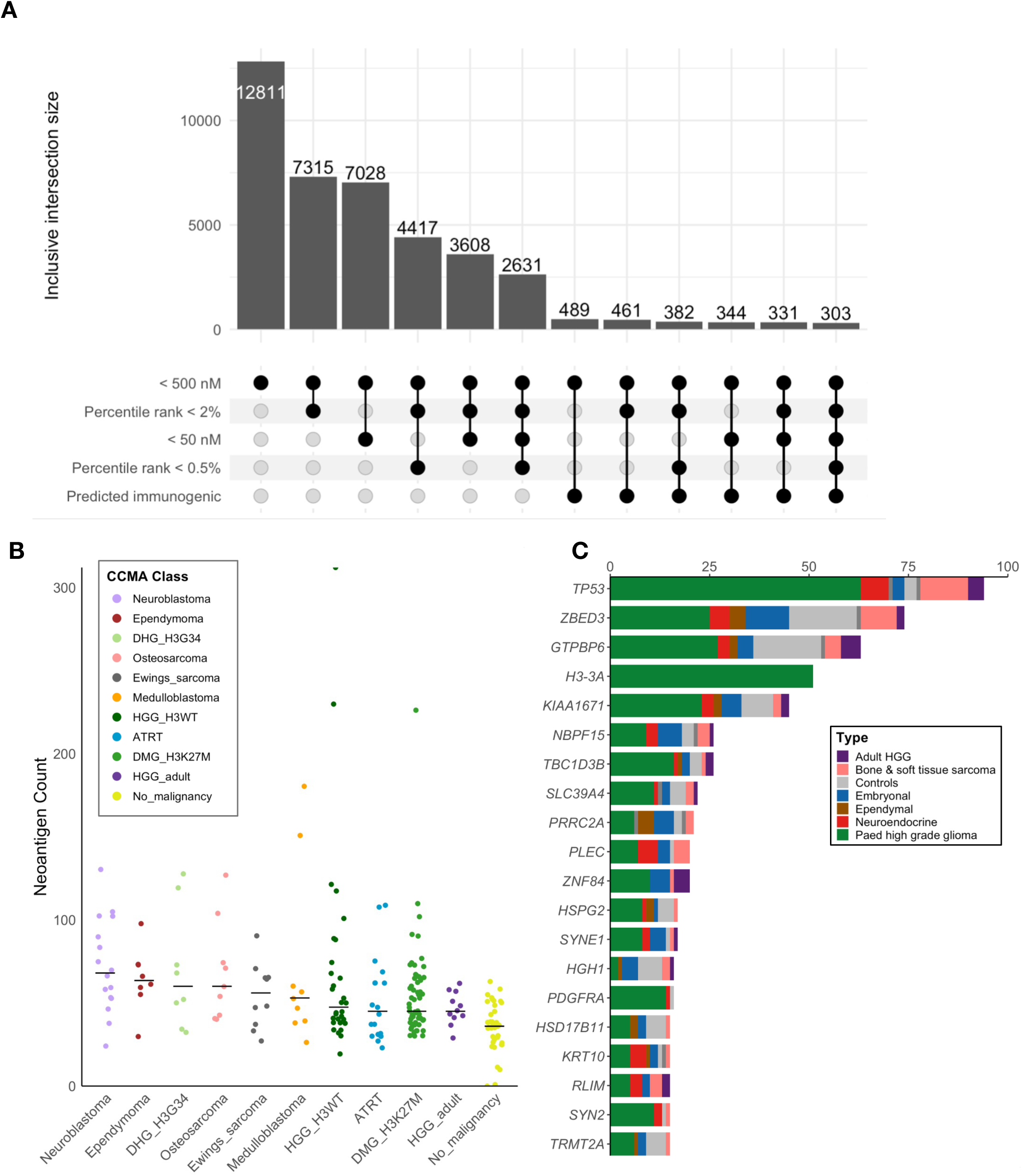
Predicted neoepitope profiles for CCMA. (A) Subgroup intersections between filtering criteria for predicted neoepitopes. Thresholds were determined by three types of prediction algorithms favoring different epitope qualities (binding, elution, and immunogenicity models). For binding and elution criteria, weak binders (IC50 <500 nM; percentile ranking <2%) and strong binders (IC50 <50 nM; percentile ranking <0.5%) were considered. For immunogenicity, a threshold score of 0.5 was considered. (B) Distribution of predicted neoepitopes with IC50 <500 nM across 11 CCMA cancer classes. The median number of neoepitopes for each class was labeled. (C) Top 15 recurrently mutated genes within the predicted neoepitope catalog. Distribution of the genes across different cancer types was shown as labeled.

The top 20 recurrent genes from predicted neoepitopes are shown in Figure 4C. Neoepitopes were most frequently contributed from *TP53* and *H3.3*. As expected, *H3.3* predicted neoepitopes were exclusively observed in the pHGG cancer type. The H3.3K27M mutation was the major contributor of predicted neoepitopes, followed by H3.3G34R and G34V. *TP53* predicted neoepitopes were observed across all cancer types except ependymal models. No dominant contributing variation was observed as diverse mutations resulted in distinct and unique predicted neoepitopes. Other known recurrent cancer genes with ≥ 10 occurrences include *PLEC*, *PDGFRA*, *NF1*, *ARID1A*, *BCOR*, *ACVR1*. These mutated genes were highly penetrant across cancer types with predicted effective, suggesting their potential as therapeutic targets.

To support our neoepitope prediction approach, we cross-referenced predicted neoepitopes from the CCMA with validated neoepitopes from the Immune Epitope Database^27^ (IEDB) and published literature. Ten peptides with various HLA restrictions and T-cell assays were found in the predicted neoepitopes of 13 cell line models (Table 1, Table S3.2). Validated targets included widely recognized cancer antigens (e.g., TP53, KRAS, NRAS, EGFR). Among these, the hallmark mutation of DMG, H3-3A K27M, and its derived neoepitope H3.3K27M_26-35_ (RMSAPSTGGV), were predicted to present in seven models. This neoepitope has been the subject of significant yet controversial immunogenicity research, underscoring the clinical relevance of these models for validation of presence using immunopeptidomics approach and functional studies^28–34^. It is also predicted that 6 neoepitopes from different *TP53* mutations present in 9 cell line models. Several neoepitope and HLA allele combinations from CCMA neoepitope predictions were validated with MHC ligand assays. Overall, the CCMA neoepitope catalogue provides an ideal cohort of preclinical models, serving as a foundation for hypothesis-driven studies of neoantigen-targeted T-cell therapies in pediatric cancers.

**Table 1.**
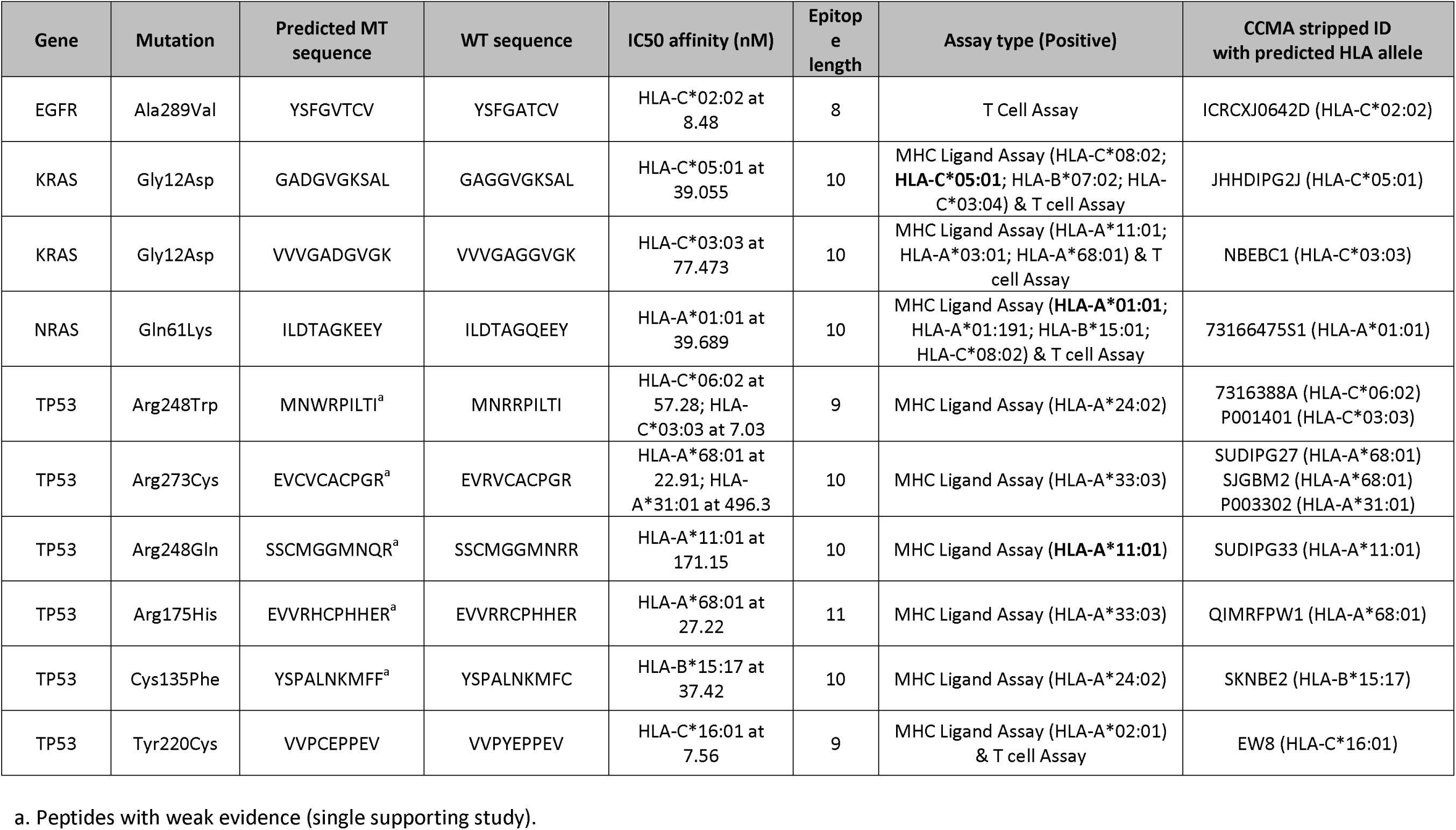
Peptides from IEDB labelled “neoepitopes” predicted in the CCMA cohort. Validated HLA restrictions matched with predicted binding of HLA allele was shown in bold.

### Immunogenicity of the CCMA Cohort Models

HLA-I molecules present intracellular antigens to cytotoxic T cells, while HLA-II molecules specialize in presenting extracellular antigens to helper T cells (Figure 5A). We firstly assessed the expression level (log_2_(TPM+1)) of individual HLA-I genes (HLA-A, HLA-B, HLA-C) to uncover their intrinsic capacity for immune engagement (Figure 5B). Specifically, high-grade gliomas exhibited relatively high HLA-A (H3G34-DHG: median=9.54, H3K27M-DMG: median=9.47, H3WT-HGG: median=8.79, HGG-adult: median=9.49) and HLA-C expression (H3G34-DHG: median=8.31, H3K27M-DMG: median=7.85, H3WT-HGG: median=7.71, HGG-adult: median=8.40), which likely contributes to more effective antigen presentation and potential immune visibility in these tumor types. Conversely, embryonal tumors demonstrated lower HLA-I expression, with notable variability among HLA-A (ATRT: median=7.21, ETMR: median=6.86, MRT: median=5.99, medulloblastoma: median=6.77), HLA-B (ATRT: median=6.46, ETMR: median=6.17, MRT: median=5.53, medulloblastoma: median=3.92), and HLA-C (ATRT: median=7.97, ETMR: median=6.36, MRT: median=6.79, medulloblastoma: median=5.87), suggesting a limited and less uniform capacity for immune engagement. Additionally, given the limited immune stimuli in the tissue culture environment, HLA-II expression (overall median value 0.1), such as HLA-DQ and HLA-DR, were generally lower in the CCMA cohort compared to HLA-I genes (overall median value 8.25; Figure S2A).

**Figure 5.**
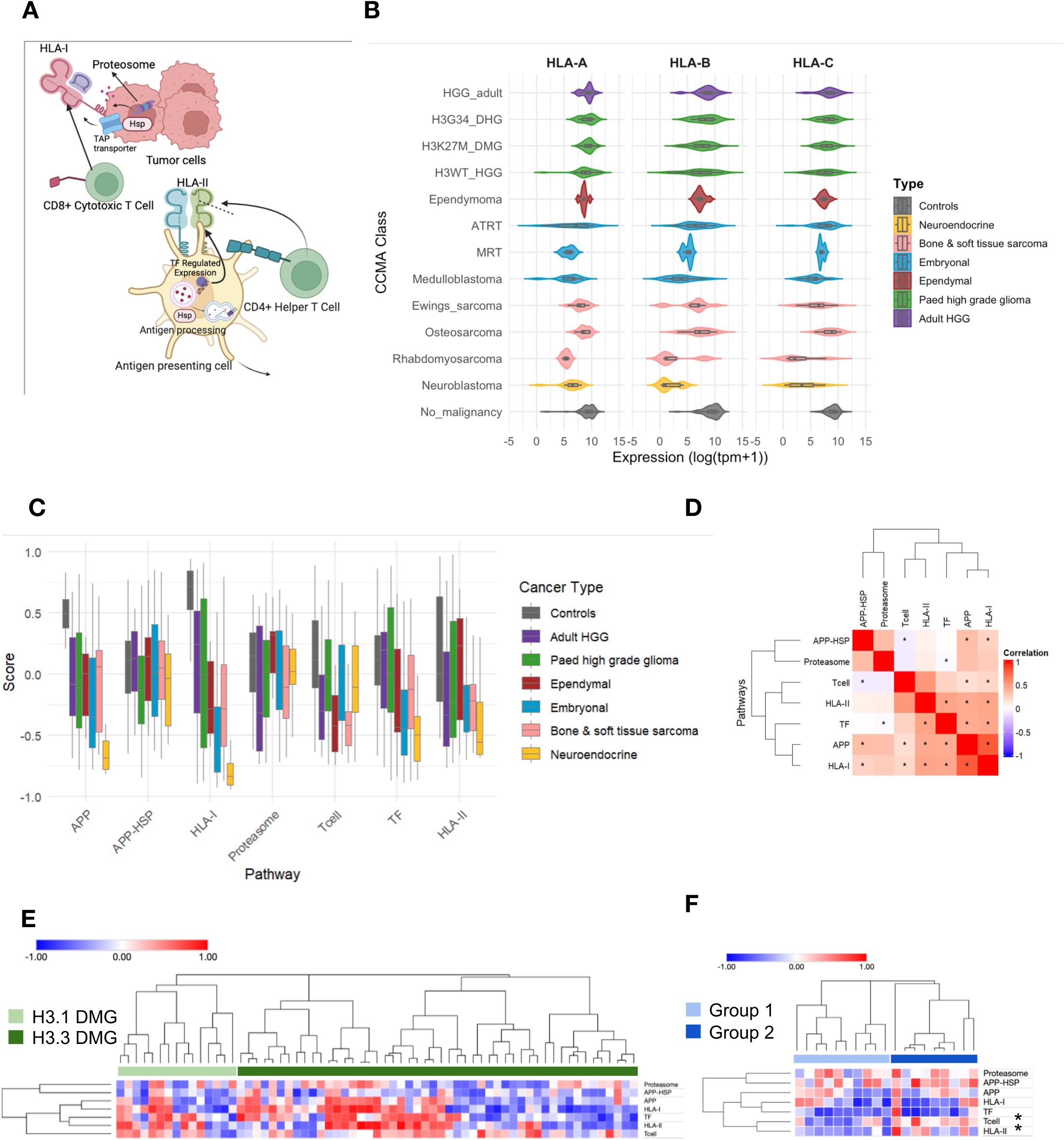
Characterization of immunogenicity of CCMA cohort using RNA-seq data. A) Schematic plot showing immunogenicity cellular pathways and key regulators. B) Violin plots showing the expression of individual HLA genes (HLA-A, HLA-B, HLA-C) across CCMA cancer types. No significant differences were identified using one-way ANOVA. C) Aggregated pathway boxplot categorized by cancer types, highlighting differential immune-related pathway activity. D) Correlation plot illustrating significant relationships among APP, HLA-I, HLA-II, and proteasome pathways. E–F) Heatmaps showing pathway activity scores for key immune-related and proteasome pathways across distinct subtypes of H3K27M-DMG (E) and ATRT (F), namely DMG-H3.1K27M and DMG-H3.3K27M, Group 1 and Group 2 ATRT. Each row represents a specific pathway, and each column corresponds to an individual tumor sample. Color scheme indicates relative expression levels, red indicates increased activity, white neutral expression, and blue downregulation. Hierarchical clustering, using Ward’s method with Euclidean distance. * Denotes FDR-adjusted p <0.05 assessed using a two-tailed Student’s t-test or Wilcoxon rank-sum test, with multiple testing correction via the Benjamini-Hochberg procedure.

In addition, we assessed the activity of key pathways that regulate antigen presentation and immune response, which are critical for understanding tumor immunogenicity and identifying vulnerabilities in immune-evasive tumors (Figure 5A). The proteasome degrades intracellular proteins into peptides, which are then processed and loaded onto MHC class I molecules via the antigen processing and presentation (APP) pathway. APP-HSP, involving HSP70 and HSP90 heat shock proteins, facilitates peptide stabilization and transport, enhancing immune recognition. Key transcription factors (TFs), such as CREB1, CIITA, and ICAM1, regulate the expression of immune-related genes, including those involved in antigen processing and HLA-I/II expression, thereby influencing tumor antigen presentation. Additionally, T-cell-related pathways modulate activation and differentiation, further shaping the immune response. The coordinated expression of HLA-I/II determines tumor visibility, directly impacting immune recognition. In CCMA, embryonal tumors exhibited the lowest immunogenicity levels (Figure 5C; consistent with the low expression of HLA genes Figure S2B). Whereas neuroblastomas showed high immunogenicity, reflected in elevated activity of the proteasome, APP, and APP-HSP pathways. Conversely, bone & soft tissue sarcomas (Ewing’s sarcoma, osteosarcoma, and rhabdomyosarcoma) demonstrated low APP activity, indicative of immune evasion mechanisms. Interestingly, tumor-associated cells (Controls) within the tumor microenvironment, such as tumor fibroblasts, exhibited stronger expression of APP, APP-HSP, HLA-I and HLA-II, suggesting that the microenvironment may play a compensatory role in antigen presentation. Notably, HLA-I gene sets showed the most variable expression patterns, reflecting fundamental differences in how tumors present peptides on the cell surface. Furthermore, we conducted correlation analyses (Figure 5E) to reveal potential co-regulation of the pathways. These analyses revealed a significant positive correlation between APP and proteasome activity (R=0.45, p-value <0.01), indicating that these processes may be tightly regulated together. APP activity showed strong associations not only with the proteasome but also with APP-HSP, HLA-I, and HLA-II expression, suggesting coordinated regulation within the antigen processing and presentation machinery.

CCMA holds the largest collection of high-grade brain cancer cell models, providing molecular diversity that enables deeper insights into the biology of pediatric CNS cancer subtypes. To address whether immunogenicity patterns correlate with subtype-specificity, we compared pediatric DMG with H3.3K27M (encoded by H3-3A, n=50) and H3.1 K27M (encoded by H3-C2 and H3-C1, n=15). No significant differences were identified between H3.3 with H3.1 DMG models (n=65; Figure 5E, Figure S3A) with substantial variation identified within H3.3 subgroups. These results indicate that immunogenicity has weak association with the specific H3-K27M mutation. In contrast, ATRT Group1 and Group2 exhibited clear differences in pathway activity (Figure 5F). For example, ATRT-Group 2 displayed significantly higher expression of HLA II pathway activity with fold change of 2.2 (p=0.026, FDR=0.093) suggesting a greater degree of immune recognition and potential immunogenicity compared to Group 1 (Figure 5F; Figure S3B). T cell signaling pathways also exhibited a clear subtype specific trend, with ATRT-Group 1 showing notably lower activation (p=0.010, FDR=0.081) (Table S4). This difference in T cell pathway activity indicates a more immunosuppressed microenvironment in Group 1 relative to Group 2, pointing to distinct immunological profiles that may influence therapeutic responses.

### CCMA Models Maintain Comparable HLA-I Expression Levels to Matched Tumors

A central question is whether the histology-specific hierarchy of HLA Class I (HLA-A, HLA-B, HLA-C) expression observed in the CCMA cohort is preserved in primary tumors. Building upon the immunogenic profiles established in the CCMA cell lines, we next compared their transcriptomic data to pediatric primary tumors and additional pediatric cancer–derived cell lines (PedCan) from the Open Pediatric Cancer (OpenPedCan) initiative. This comprehensive dataset comprises 335 solid tumor samples and 155 cell lines covering more than 20 pediatric cancer histology, providing a robust framework for assessing whether *in vitro* models faithfully recapitulate *in vivo* tumor biology^35,36^.

Given that cell lines lack tumor-intrinsic immune suppression^37^, higher absolute expression levels of HLA-I genes might be expected *in vitro*. A two-way ANOVA test (p <0.001) (Figure S4 and Table S5) confirmed significant differences between sample types, i.e. cell lines and tumor tissues, but the preservation of cancer-specific patterns underscores the intrinsic nature of these immunogenic profiles (Figure 6). Indeed, we found that the relative ordering of tumor subtypes largely remains comparable: neuroblastomas and ependymomas exhibit higher HLA Class I expression, whereas ATRT and medulloblastoma rank lower (Figure 6). In contrast to HLA Class I, HLA Class II genes (e.g., HLA-DQ, HLA-DR) were significantly more highly expressed in primary tumors than in any of the cell line cohorts, due to lack of immune stimuli in the tissue culture system (CCMA or PedCan; Figure 6 A-C; Figure S4). This observed upregulation of HLA Class II gene expression in primary tumors, compared to cell line cohorts, underscores the pivotal role of the tumor microenvironment-encompassing cytokines, immune infiltrates, and other regulatory signals-in modulating these immune-related processes. Beyond individual HLA genes, we examined whether the proteasome and antigen processing and presentation (APP) pathways exhibit histology specific patterns in both primary tumors and cell lines (Figure S5). Despite the presence of cytokine driven modulation *in vivo*, key pathway trends identified in CCMA derived cell lines were largely mirrored in solid tumor samples. In particular, neuroblastoma showed significantly elevated proteasome and APP activity (p < 2 × 10^−14^ for cancer subtype, p < 8 × 10^−21^ for sample type, p < 4 × 10^−8^ for interaction), a pattern coinciding with its higher HLA Class I expression. By contrast, ATRT remained consistently low across sample types, reflecting a more immune-evasive phenotype. These highly significant two-way ANOVA findings (Figure S5) indicate that while many core antigen-processing and presentation mechanisms are conserved, their relative activity can shift depending on tumor subtype and *in vitro* versus *in vivo* contexts.

**Figure 6.**
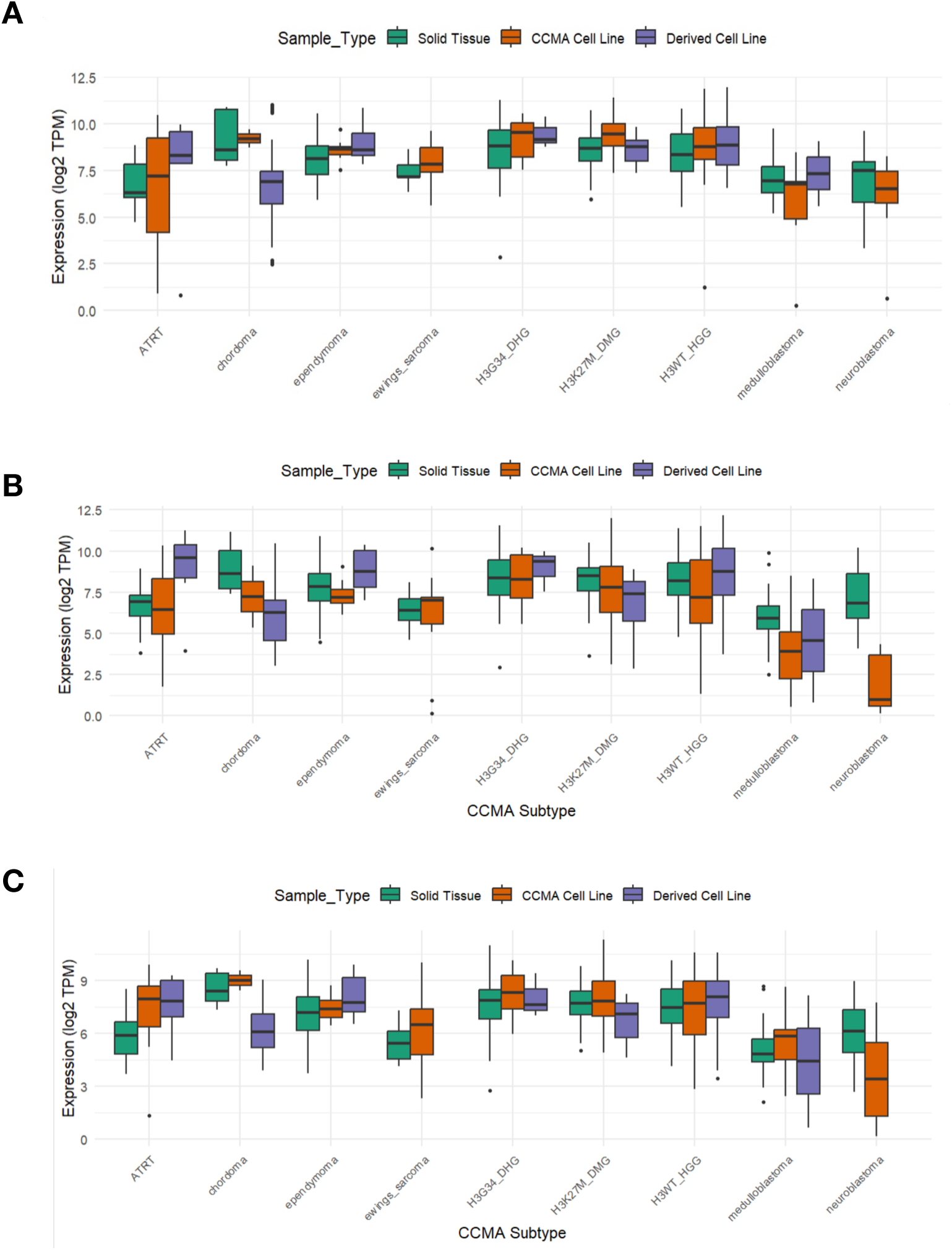
HLA Class I Gene Expression Across Pediatric Cancer Subtypes in Different Sample Types. (A–C) Box plots displaying log2-transformed transcript per million (TPM) expression values of HLA-A (A), HLA-B (B), and HLA-C (C) across various pediatric cancer subtypes in solid tumor samples (green) and derived cell lines (purple) from OpenPedCan, and CCMA cell lines (orange) from our analysis.Statistical significance was determined by two-way ANOVA test (p <0.001).

## Discussion

While extensive immunogenic characterizations and HLA typing has been reported for large cancer cell line cohorts such as CCLE^38^, pediatric cancer models have lagged behind these efforts. HLA typing is essential for preclinical research, particularly in advancing immune-based therapies, including personalized vaccines, TCR-T therapies, and other immunotherapeutic approaches. In this study, we compiled a comprehensive catalog of Class I HLA types and neoepitopes for nearly 300 high-grade pediatric cancer cell lines in the CCMA, which includes both widely used and newly established models. This dataset represents the largest collection of RNA-seq data for pediatric high-grade cancer cell models, particularly CNS tumors. This publicly available resource serves as a centralized dataset, enabling researchers to accelerate hypothesis testing and streamline preclinical investigations. We analyzed the immunogenic landscape, assessing HLA-I and HLA-II expression across different cancer types and subtypes.

The CCMA demonstrated broad coverage of HLA alleles and supertypes, including prevalent alleles across ethnicities^39^ and rare alleles with dual specificity supertypes (“A01 A03” and “A01 A24”). Previous studies have investigated associations between HLA supertypes and clinical outcomes in adult cancers. The B44 supertype was reported to be associated with extended survival in melanoma patients with ICB therapy. Whereas the B62 supertype was associated with poor outcome^16^. Less information is known regarding HLA supertypes in pediatric cancers. Few studies showed evidence of HLA-DPB1 supertypes providing a protective role in childhood leukemia. A negative association was observed between the DPB1 supertype with cancer prognosis^40^. Our atlas provides a well-balanced pediatric cancer cohort for immunotherapy studies and is suitable for informing study design with various HLA allotype and supertype restrictions.

The CCMA encompasses models with varying degrees of HLA-I homozygosity, with 23 models featuring homozygous HLA types across all HLA-I genes (<u>Table S1.1</u>). We suspect the high homozygosity frequency stems from HLA LOH through aberrations to chromosome 6 or focal deletions of the HLA locus. Furthermore, HLA-I allele-specific transcriptional downregulation was detected in 24 models. Both observations lead to HLA LOH which restricts peptide presentation and, therefore, are important factors to consider when selecting cancer models for testing hypotheses. Disrupted HLA-I expression helps tumors evade immune detection, but complete loss increases their vulnerability to NK cell surveillance due to the absence of self-antigen presentation^20^. Therefore, partial HLA-I down-regulation may enable tumors to escape both T-cell and NK surveillance^41^. HLA LOH and ASE loss have been frequently observed across many adult cancer types, including pancreatic cancer^42^, bladder cancer^43^, and lung cancer^44^. HLA LOH can also occur in pediatric cancer^45^. Consistent with our findings, HLA homozygosity and LOH were more frequently observed in sarcomas, particularly osteosarcomas^45^, which is likely associated with the known genomic instability. The HLA-I homozygous CCMA models provide a valuable resource to elucidate the ramifications of a reduced epitope repertoire and HLA LOH, whereas cell line models exhibiting reversible ASE loss can be used to investigate the therapeutic benefit in recovering expression of the lost HLA allele with interferons or inhibitors^46,47^. However, extensive passaging of cell lines can introduce excess genetic stress and contribute to LOH^48^. Further validation with tools such as LOHHLA^18^ or DASH^19^ for HLA LOH and arcasHLA-quant^22^ for ASE loss is required to determine whether HLA LOH in CCMA models is a tumor-driven observation.

*sIn silico* prediction of neoepitopes from CCMA models identified several public neoantigens derived from recurrent cancer driver mutations (Table 1, Table S3.2). We found H3.3K27M derived neoepitopes predicted to bind to various HLA alleles in patient models. The H3.3K27M mutation is highly recurrent in pediatric HGG. The HLA-A*02:01 restricted neoepitope (RMSAPSTGGV) has shown promise in T cell recognition^28,29^ and its peptide vaccine resulted in increased survival in pediatric DMG patients^30^. However, recent evidence suggests the T cell recognition of the RMSAPSTGGV neoepitope cannot be replicated and is not endogenously presented in DMG models^31,34^. Despite this, the H3.3K27M mutation remains a prime neoantigen target due to its recurrence and broad applicability to DMG patients. One putative H3.3K27M neoepitope predicted from our study was the ATKAARMSA peptide, which was predicted to bind to HLA-A*30:01 with high affinity and high likelihood of presentation (percentile rank of 0.02 by netMHCpanEL) in five models. Notably, the ATKAARMSA peptide was also predicted from the study by Chang et al., 2017^49^ with NetMHCcons v1.1. The H3.3K27M22-30 (ATKAARMSA) peptide may be a promising candidate for immunogenicity validation. Furthermore, growing evidence suggests the importance of HLA-II restricted antigen presentation in cancer immunotherapies^50,51^. The INTERCEPT H3 trial utilizes a long H3.3K27M_14-40_ peptide to elicit CD4^+^ T cell-mediated immune responses without HLA-A*02:01 restrictions^52^.

Other hallmark pediatric cancer mutations including the SWI/SNF subunit SMARCB1 frameshift/termination/deletions for ATRT/MRT and MYC duplications for medulloblastoma were not predicted as effective neoepitopes. This was primarily limited by the type of mutations considered in this study. *TP53* mutations and predicted neoepitopes were also frequently found in CCMA models. The TP53Y220C_217-225_ (VVPCEPPEV) neoepitope has been well characterized for immunogenicity from multiple studies^53,54^ (Table 1, Table S3.2). Notably, CD8^+^ T cell responses to the TP53R248W_240-249_ (SSCMGGMNWR) neoepitope restricted to HLA-A*68-01 was observed in patients with epithelial cancers^53,55^. Although the SSCMGGMNWR neoepitope was not predicted from our data, we observed predicted binding and presentation (percentile rank of 0.49 by netMHCpanEL) from the TP53R248Q_240-249_ (SSCMGGMNQR) peptide with HLA-A*11:01 in the SUDIPG33 model (Table 1, Table 3.2). Interestingly, the well-studied TP53R175H_168-176_ (HMTEVVRHC) neoepitope^56^ was not identified from our predictions. As previously reported, the HMTEVVRHC peptide performs poorly in computational binding prediction^56,57^. While we found the TP53R175H mutation present in four HLA-A*02:01^+^ cell lines, the HMTEVVRHC neoepitope was predicted by NetMHCpan 4.1 as a non-binder to HLA-A*02:01 at 4543.17 nM (rank 7.5%). These cases underscore the importance of understanding the propensity for Type I & II errors that inevitably result from all current neoepitope prediction algorithms. With this in mind, the demonstrable strengths and benefits of current neoepitope prediction approaches provide valuable utility and enable candidate discovery and prioritization for downstream experimental validation. The CCMA provides a valuable platform to investigate peptide processing, presentation, and immunogenicity for novel HLA allele and neoepitope combinations.

CCMA cell lines largely preserved histology-specific HLA-I expression patterns, supporting their relevance for immuno-oncology research. However, primary tumors exhibited higher HLA-II expression, likely driven by immune-stimulating factors such as cytokines and interferons present *in vivo*. These findings suggest that tumor models can effectively capture intrinsic antigen presentation dynamics but may require additional microenvironmental context to fully replicate immune interactions.

Our comprehensive evaluation of immunogenicity in the CCMA cohort underscores the varying degrees of immune engagement across pediatric tumor types and subtypes and highlights the influence of both intrinsic tumor characteristics and extrinsic microenvironmental factors. High-grade gliomas (e.g., H3K27M-DMG, H3G34-DHG) exhibited elevated HLA-I (HLA-A, HLA-B, HLA-C) expression, facilitating stronger immune recognition, while embryonal tumors (e.g., medulloblastoma, MRT) displayed significantly lower levels, suggesting an immune-cold phenotype with reduced antigen presentation potential. Cellular pathway analysis reinforced these findings, revealing that embryonal tumors consistently showed lower proteasome and APP activity, indicative of an intrinsic tendency toward immune evasion. These findings align with earlier reports demonstrating that HLA I-mediated antigen presentation can significantly dictate tumor visibility to cytotoxic T cells^58,59^. Further analysis revealed co-regulation between proteasome and APP pathways, particularly emphasizing the role of heat shock proteins (e.g., HSP70, HSP90) in stabilizing peptides and facilitating MHC class I loading^60^. Notably, subgroup-level heterogeneity was evident, with ATRT Group 2 (MYC-high) demonstrating higher HLA-II expression and greater T-cell signaling activity than Group 1, underscoring molecular subgroup-specific differences in immune engagement^61^. While this distinction indicates that Group 2 tumors may have a greater capacity for immune engagement, ATRT as a whole remains characterized by low baseline immunogenicity and immune evasion mechanisms.

Therapeutically, our findings indicate that tumors with robust antigen-processing activity (e.g., high-grade gliomas) may benefit from interventions that further enhance antigen presentation, such as interferon-γ–based treatments or proteasome modulators. Conversely, immune-cold tumors like medulloblastoma and ATRT may require alternative strategies, including epigenetic modifications to upregulate MHC expression or adoptive cellular therapies to bypass conventional antigen presentation constraints. Recognizing these tumor- and subgroup-specific immunogenic signatures could refine immunotherapeutic approaches, enabling more targeted and effective treatments for pediatric malignancies.

The characterization of immunogenic features reported in this study carries significant clinical implications, especially given the low survival rates and considerable treatment-related toxicities experienced by pediatric cancer patients. This cohort prominently features CNS tumors, which remain the leading cause of disease-related mortality, while also including rare and under-characterized pediatric cancer types. Our resource paves the way for novel immunotherapeutic strategies aimed at pediatric cancers associated with poor survival outcomes and severe long-term side effects. Although the current study primarily emphasizes HLA class I typing and neoepitope prediction in high-risk tumors, it also provides a versatile platform that can be expanded to incorporate additional molecular characteristics relevant for advancing immunotherapy development

## Limitations of the Study

While this comprehensive analysis advances our understanding of pediatric cancer immunogenicity, several limitations warrant consideration. First, neoepitope predictions are based on computational algorithms and are subject to both false positives and false negatives, underscoring the need for experimental validation. Second, prolonged passaging of cell lines across different laboratories may introduce genetic drift and varying selective pressures, including HLA loss of heterozygosity, potentially confounding interpretations of HLA homozygosity and allele-specific expression loss. Third, the absence of the native tumor microenvironment in cell lines limits the extrapolation of these findings to *in vivo* contexts. Despite these limitations, the resulting atlas offers a valuable resource to support hypothesis generation and prioritization for downstream experimental and translational studies in pediatric cancer immunotherapy.

## Resource availability

### Lead Contact

Further information and requests for resources and reagents should be directed to and will be fulfilled by the Lead Contact, Dr Claire Sun.

### Materials availability

This paper does not report novel material or reagents generated.

### Data and code availability

- Data sets are publicly available at vicpcc.org.au/dashboard and deposited at “CCMA primary multi-omics datasets”, Mendeley Data, V1, doi: 10.17632/rnfs539pfw.1. HLA typing raw output from WGS and RNA sequencing are in Table S1.
- This paper does not report original code.
- Any additional information required to reanalyze the data reported in this paper is available from the lead contact upon request.

## Acknowledgments

Funding for the project is from the Medical Research Future Fund MRF2030828. This project is also supported by the Next generation Precision Program which is/was generously supported by the Children’s Cancer Foundation Australia, Robert Connor Dawes Foundation, and the Medical Research Future Fund (NHMRC Project 2007620). C.S. is funded by Australia Victoria Cancer Agency ECRF22006. We thank over 30 contributing sites for providing cell lines and/or tissues from which cell lines were generated. A full list of cell line contributors is provided in the supplemental information.

## Author contributions

*The studies were conceived and designed by C.S. and R.F.;* Y.G., I.M. and C.S. conducted computational analysis and wrote the paper; V.A. conducted the analysis for HLA calling using Optitype. I.W. contributed to analysis design and writing; T.S. and P.F. provided NGS HLA typing results.

## Methods

### Cell line sequence data

Whole genome and RNA sequences for 287 (249 and 275 respectively) cell lines were obtained from the Childhood Cancer Model Atlas as described in Sun et al., 2023^4^ (n=174) and newly updated datasets (n=113). Briefly, primary cell lines were generated in-house from tumor tissue or obtained from collaborators. Cell lines were validated by identifying hallmark genetic attributes through WGS or STR profiling.

WGS was performed on extracted genomic DNA (DNeasy Blood & Tissue Kit) with 150bp paired-end sequencing. A minimum of 30X coverage was achieved for all cell lines. Raw data was processed using the nf-core Sarek pipeline (version 3.4.4).

Briefly, sequences were preprocessed with GATK4 Best Practices and mapped to the human reference genome GRCh38 with BWA-MEM. Variants were called with Mutect2 and annotated with snpEff and Ensembl VEP. For RNA sequencing, RNA was extracted (RNeasy Kit), ploy-A enriched, processed as a sequencing library, and sequenced with paired-end 150bp sequencing. RNA expression levels were expressed as log2(TPM+1).

### NGS amplicon sequencing high-resolution HLA typing

High-resolution HLA typing with NGS amplicon sequencing was performed for 29 cell lines (23 pediatric HGG; 5 Bone & soft tissue sarcoma; 1 neuroblastoma; 1 control). Samples were processed at the Victorian Transplantation and Immunogenetics Service (West Melbourne, Victoria, Australia) and PathWest Laboratory Medicine (Perth, Western Australia, Australia). Complete or partial 4-digit, 6-digit, or 8-digit classical class I and II HLA types were obtained. IMGT/HLA sequence database versions 3.38.0, 3.50.0.0, and 3.53.0.0 were used as the reference database.

### HLA typing

The nf-core hlatyping pipeline (version 2.0.0) was used to determine 4-digit class I HLA types for 287 unique cell lines. In brief, raw DNA and RNA sequences were processed with FastQC, mapped to the human reference genome GRCh38 with Yara, and HLA types were determined with Optitype. Default settings were used, with enumerations set to 3. IMGT/HLA release 3.14.0 was used as the reference HLA sequence.

Four-digit HLA types were determined for the classical class I genes (HLA-A, HLA-B and HLA-C) with a decision-making tree (Figure 1B). High-resolution HLA types were reported when available, as they are considered the most accurate method, superseding the NGS sequence-based typing results generated by Optitype.

Manual curation was used to determine non-concurrent HLA types. Cases include a match in 2-digit but not the 4-digit resolution or a complete mismatch between WGS and RNA-seq inferred typing. The alternative solutions (solutions 1 and 2) were used to resolve ambiguous HLA types. Preference was given to high-confidence alleles predicted across all three solutions. The curated HLA types were reported with an ambiguity flag. Cell lines with homozygous HLA types for all three genes were also flagged.

After determining the definitive HLA types, HLA supertypes for genes A and B were assigned to each cell line. HLA supertypes were defined according to the classification by Sidney et al., 2008^8^. Before assigning the supertypes, HLA types were converted to the new nomenclature based on HLA allele designations from April 1st, 2010. All represented HLA alleles in our dataset were covered by supertype classifications.

### Accuracy evaluation

Overlapping results between NGS high-resolution typing and Optitype were used to validate and calculate the accuracy of our strategy. Typing accuracy and zygosity prediction accuracy were defined according to Szolek et al., 2014^5^. Briefly, typing accuracy was based on the percentage of correctly predicted 2-digit or 4-digit alleles. Zygosity prediction accuracy was determined by the percentage of correctly predicted homozygous and heterozygous loci, disregarding the correctness of the typed allele.

### Neoepitope prediction

Neoepitope candidates were identified using the pVACseq cancer immunotherapy pipeline (version 4.4.1). Cell line-specific HLA types and mutation data were prepared and preprocessed according to pipeline instructions. Mutations were filtered to exclude germline variants labelled by Mutect2. Variants passing the Mutect2 hard filter were retained. The hard filter tags include clustered-events, duplicate, multiallelic, base-qual, map-qual, fragment, position, panel-of-normals, normal-artifact, and contamination. Variants were then annotated using the Variant Effect Predictor (VEP; release 113) with the Frameshift and Wildtype plugins. To reduce computational load, the “--pick” option was used to retain the most impactful transcript for each mutation.

A variant consequence filter was applied to retain variants in the coding region. Using sequence ontology (SO) terms, variants with a consequence matching any child terms of coding-sequence-variant (SO:0001580) were selected. The retained SO terms include, stop-gained, frameshift-variant, stop-lost, start-lost, inframe-insertion, inframe-deletion, missense-variant, protein-altering-variant, incomplete-terminal-codon-variant, start-retained-variant, stop-retained-variant, synonymous-variant, and coding-sequence-variant.

Gene expression data was integrated using VAtools (version 5.1.1) to annotate variants with gene expression levels in TPM. Somatic variants were used as input for pVACseq. A separate input of phased variants combining somatic and germline variations was prepared to account for proximal variants. Briefly, Mutect2 hard-filtered somatic variants and variants with the germline tag were retained. The combined variants were annotated with VEP and filtered for coding variants as described. ReadBackedPhasing in GATK (Version 3.6) was used to generate the phased variant input from WGS reads.

The pVACseq pipeline was executed to generate all possible 8, 9, 10, and 11-mer epitopes. Seven prediction algorithms were used to evaluate peptide-MHC binding affinity (NetMHCpan-BA, MHCflurry, MHCnuggets-I), elution (NetMHCpan-EL, BigMHC-EL, MHCflurryEL), and immunogenicity (BigMHC-IM). All possible downstream analysis options were specified. Epitope cleavage site was predicted with NetChop C term 3.0, peptide stability was predicted with NetMHCStabPan, and reference proteome similarity was determined with the human reference proteome GRCh38.

Promising neoepitope candidates were filtered with default parameters from pVACseq, with the IC50 scoring metric set to lowest. The applied filters include mutant allele IC50 binding scores below 500 nM, tumor DNA coverage >10, tumor DNA variant allele frequency >0.25, gene expression >1, transcript support level ≤1, and only the top epitope was reported for each variant.

### Tumor Data Sources

Gene expression data for the tumor tissues and external cell lines were obtained from the OpenPedCan repositories, which provide comprehensive RNA-sequencing profiles of pediatric cancers. The dataset included samples from key pediatric cancer subtypes, including gliomas, sarcomas, embryonal tumors, neuroendocrine and ependymal tumors. A total of 489 samples representing 9 subtypes were included after meeting the following selection criteria: (i) high RNA-seq quality scores, (ii) sufficient cancer type representation (≥ 2 samples per type), and (iii) availability of complete metadata, including histological subtype and sample type (cell line vs. solid tissue).

### Gene Set Variation Analysis

Pathway activity was assessed using Gene Set Variation Analysis (GSVA) with the GSVA R package (v4.4.0). GSVA does not require predefined sample labels, making it particularly suitable for detecting subtle and continuous changes in gene set enrichment across diverse tumor subtypes.

Pathway gene sets were curated from the Molecular Signatures Database (MSigDB) and from existing immuno-oncology literature (Table S6). Key pathways of interest included antigen processing and presentation (APP), HLA Class I and II, proteasome function, APP-HSP (heat shock protein–related antigen processing), T-cell activation, and transcription factors involved in immune regulation (e.g., CREB1, CIITA, ICAM1). Using the log_2_(TPM+1) expression matrix, GSVA calculated per-sample enrichment scores for each pathway, enabling direct comparisons across cell lines and primary tissues.

### Statistical Analysis

All statistical analyses were conducted in R (v4.4.0). Differences in pathway enrichment among subtypes were assessed using the Kruskal-Wallis test, with Dunn’s test for post hoc pairwise comparisons and Bonferroni correction to account for multiple testing. When comparing gene expression levels-such as HLA Class I genes-across both subtypes and sample types (cell lines vs. primary tumors), two-way ANOVA was applied. Post hoc tests (e.g., Tukey’s HSD) were then performed when significant main effects or interactions were detected. Correlation analyses between pathways (e.g., proteasome activity vs. APP) were performed using Pearson’s correlation coefficient. p-values from multiple comparisons were adjusted using the Benjamini-Hochberg method, and correlations meeting the adjusted threshold (FDR <0.05) were deemed significant.

### Reproducibility and Software

Key R packages included ggplot2, ComplexHeatmap, and gridExtra for data visualization, alongside the base hclust function for hierarchical clustering. All scripts, configurations, and documentation are available upon request or in the supplementary materials, ensuring transparency and reproducibility of the computational analyses.

### Data Availability

All processed RNA-seq datasets from OpenPedCan are publicly accessible through their respective repositories. CCMA RNA and SNP data are publicly accessible through its data dashboard and data portal at vicpcc.org.au/dashboard. Raw sequencing data has been published and archived at European Genome Archive EGA: EGAS00001006320 and is available from the lead contact upon agreement of the data access policy. Detailed metadata, including histological classifications and RNA expression profiles for OpenPedCan study, can be found in the Table S7 and additional technical sequencing metrics may be available through the corresponding analysis pipelines on the OpenPedCan GitHub repository. HLA typing raw output from WGS and RNA sequencing are in Table S1.1 and at Mendeley Data: https://doi.org/10.17632/rnfs539pfw.1.

## Supplementary Figure Legends

**Figure S1.**
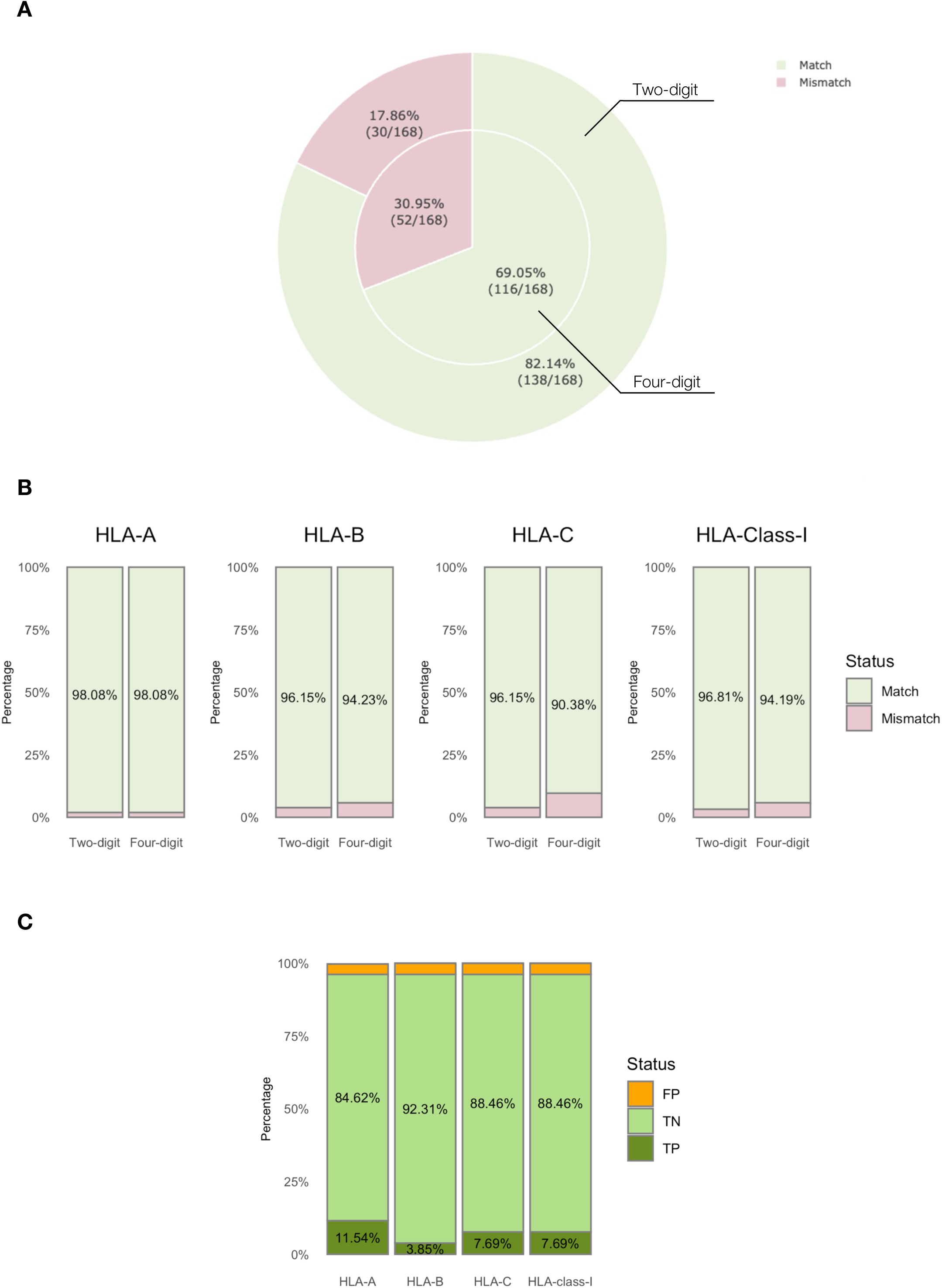
HLA typing accuracy assessment for CCMA cohort. (A) Stacked pie charts showing 2-digit and 4-digit concordance rate between mismatched cell line models. A total of 28 models with 168 HLA alleles were assessed. (B) Bar chart showing the accuracy of predicted HLA types. The percentage of match and mismatch HLA calls was inferred from comparing Optitype with high resolution typing. (C) Bar chart showing the accuracy of HLA zygosity prediction. The percentage of true positive (TP), true negative (TN), false positive (FP), and false negative (FN) classifications was inferred from comparing Optitype with high resolution typing.

**Figure S2.**
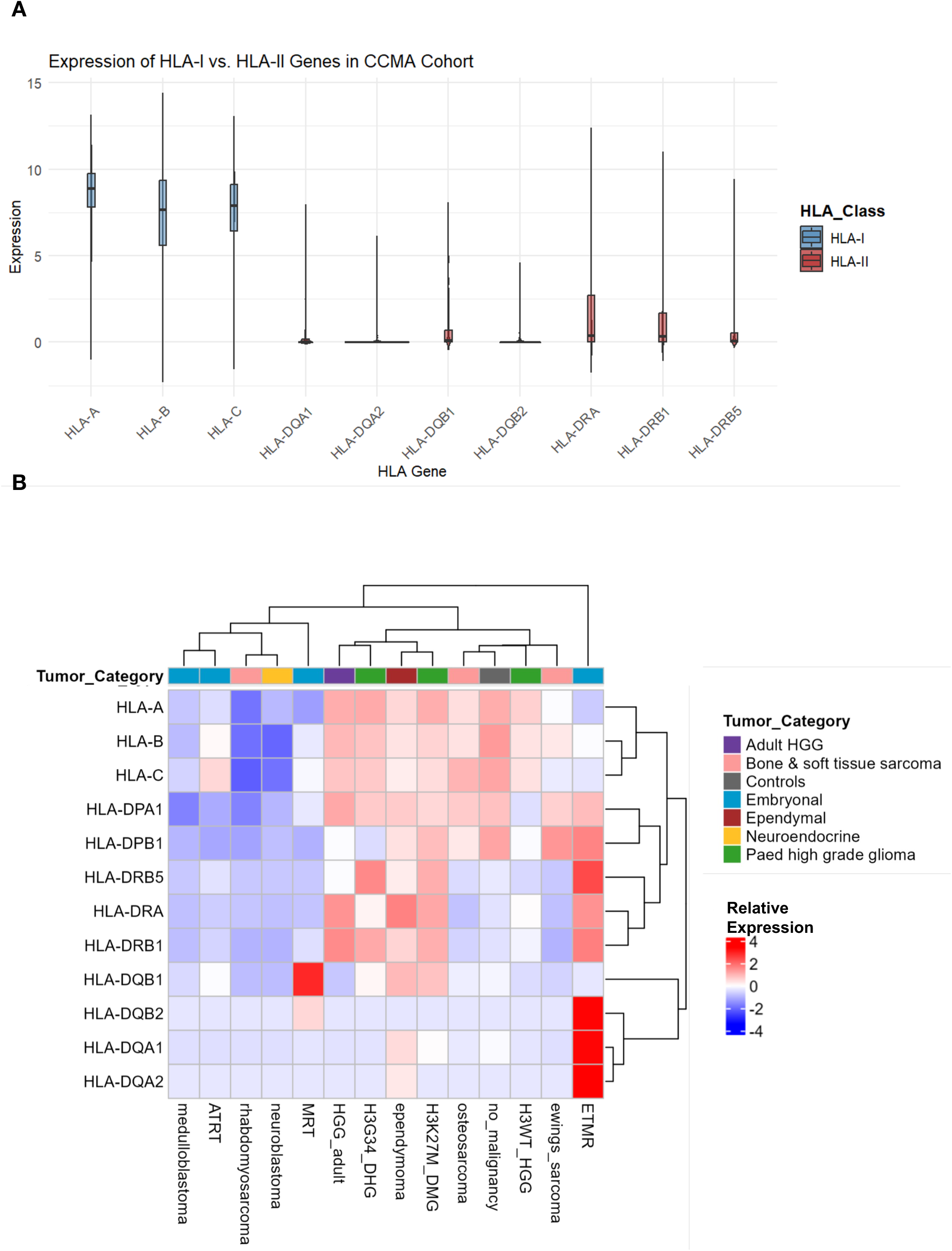
HLA Gene Expression Profiles in the CCMA Cohort. (A) Box plot comparing HLA Class I (HLA-I (blue): HLA-A, HLA-B, HLA-C) and HLA Class II (HLA-II(red): HLA-DPA1, HLA-DPB1, HLA-DQA1, HLA-DQB1, HLA-DQB2, HLA-DRA, HLA-DRB1, HLA-DRB5) gene expression (log2(TPM+1)) across tumor subtypes in the CCMA cohort. B) Heatmap showing the median expression and unsupervised hierarchical clustering of HLA-I and II genes for different cancer types in CCMA. The color scale represents per-gene z-score normalization, where red indicates higher relative expression and blue denotes lower expression. Tumor categories are annotated on the left.

**Figure S3.**
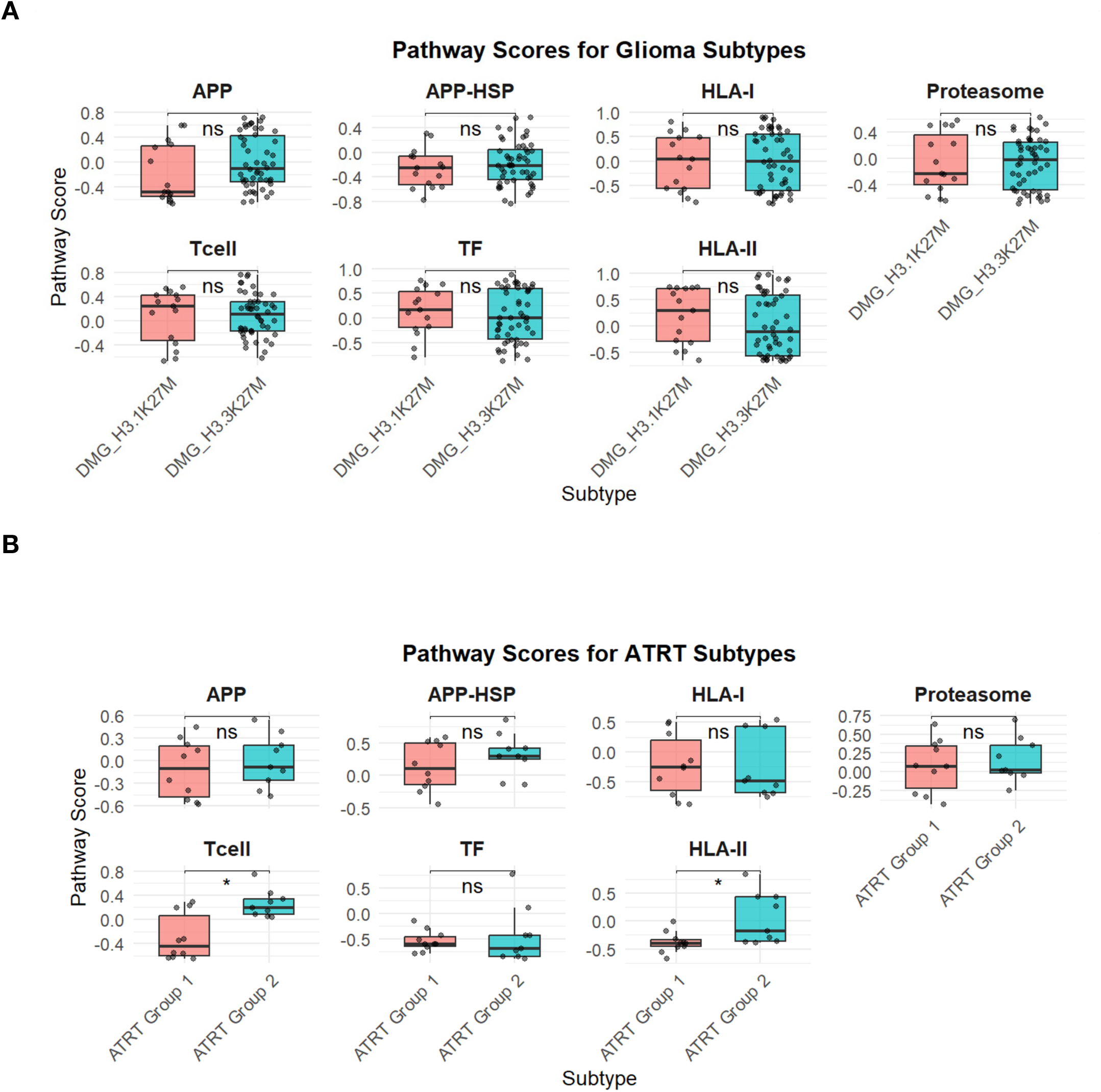
Immunogenicity pathway activity comparison between molecularly defined cancer subtypes in H3K27M-DMG (A) and ATRT (B). Student t test was used to determine the statistical difference between the pathway scores of DMG_H3.1K27M and DMG_H3.3K27M (A), ATRT Group1 and Group2 (B). Statistical significance is indicated (*p <0.05).

**Figure S4.**
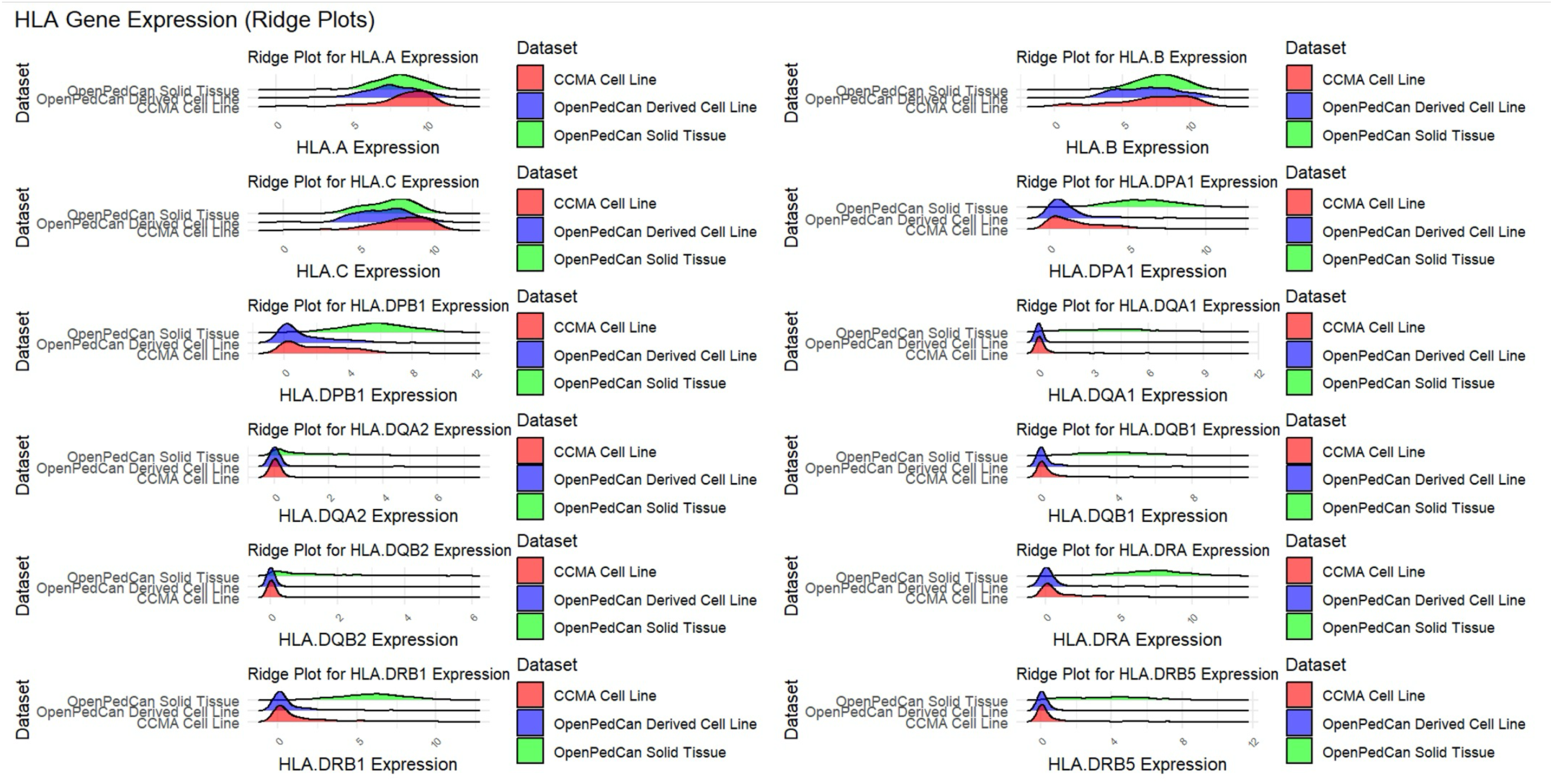
Comparison of HLA gene expression between tumor tissues and cell line models. Expression of HLA-I (HLA-A, HLA-B, HLA-C) and HLA-II (HLA-DP, HLA-DQ, HLA-DR subunits) genes are shown in ridge plots between CCMA cohort (red), openPedCan tumor tissues (green) and derived cancer cell lines (blue).

**Figure S5.**
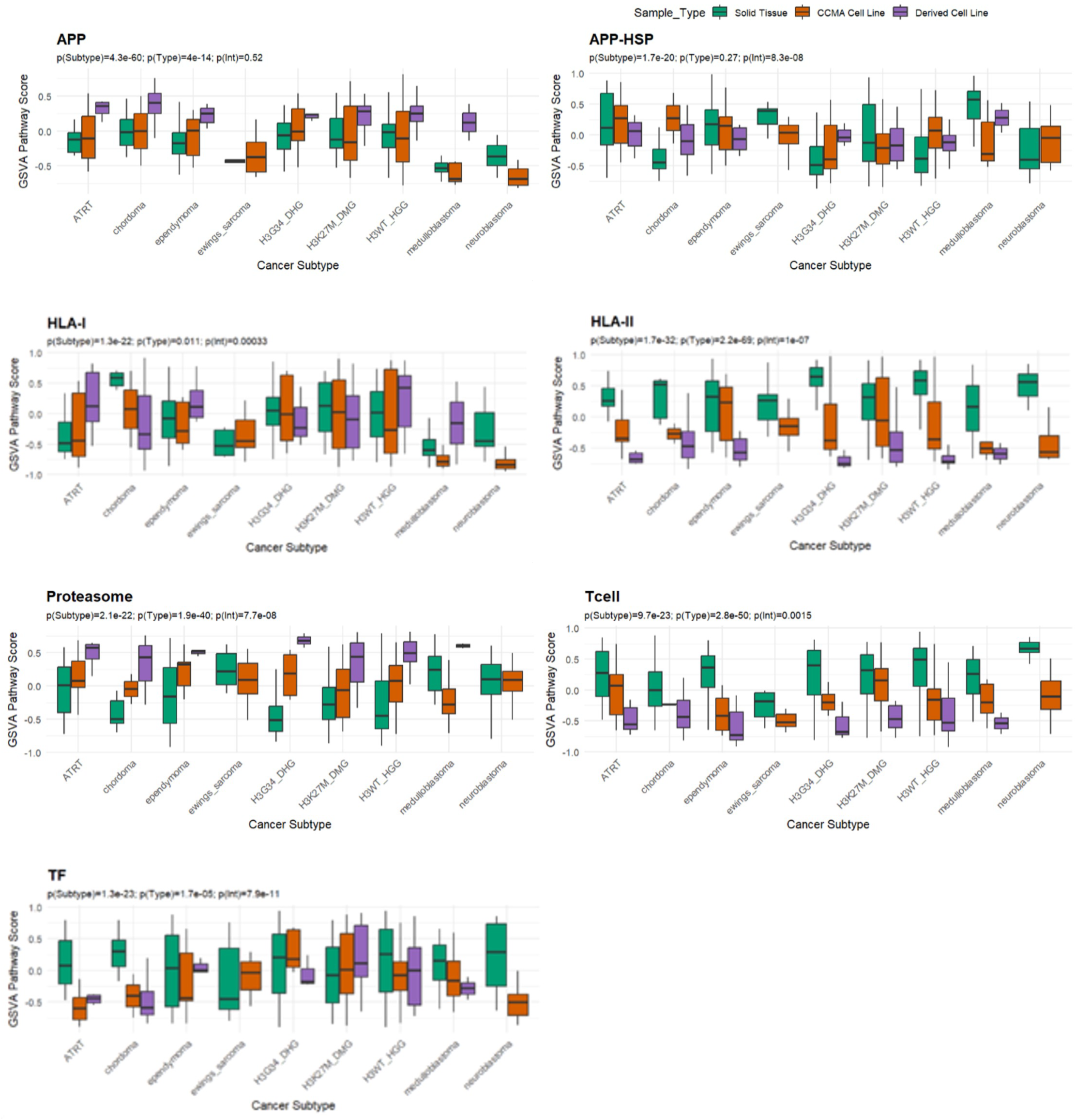
Comparison of Pathway Activity Across Cancer Subtypes and Sample Types. Box plots displaying GSVA pathway activity scores for APP, APP-HSP, HLA-I, proteasome, T cell, and TF pathways across pediatric cancer subtypes in CCMA cell lines (orange), OpenPedCan-derived cell lines (purple), and OpenPedCan solid tumor samples (green). Statistical significance from two-way ANOVA is indicated in each panel, assessing differences in pathway activity across cancer subtypes and sample types.

