## Supplementary figures and images for "Resource: A Compendium of HLA-Type and Expression in Pediatric Cancer Models"

### SupplementaryFiles

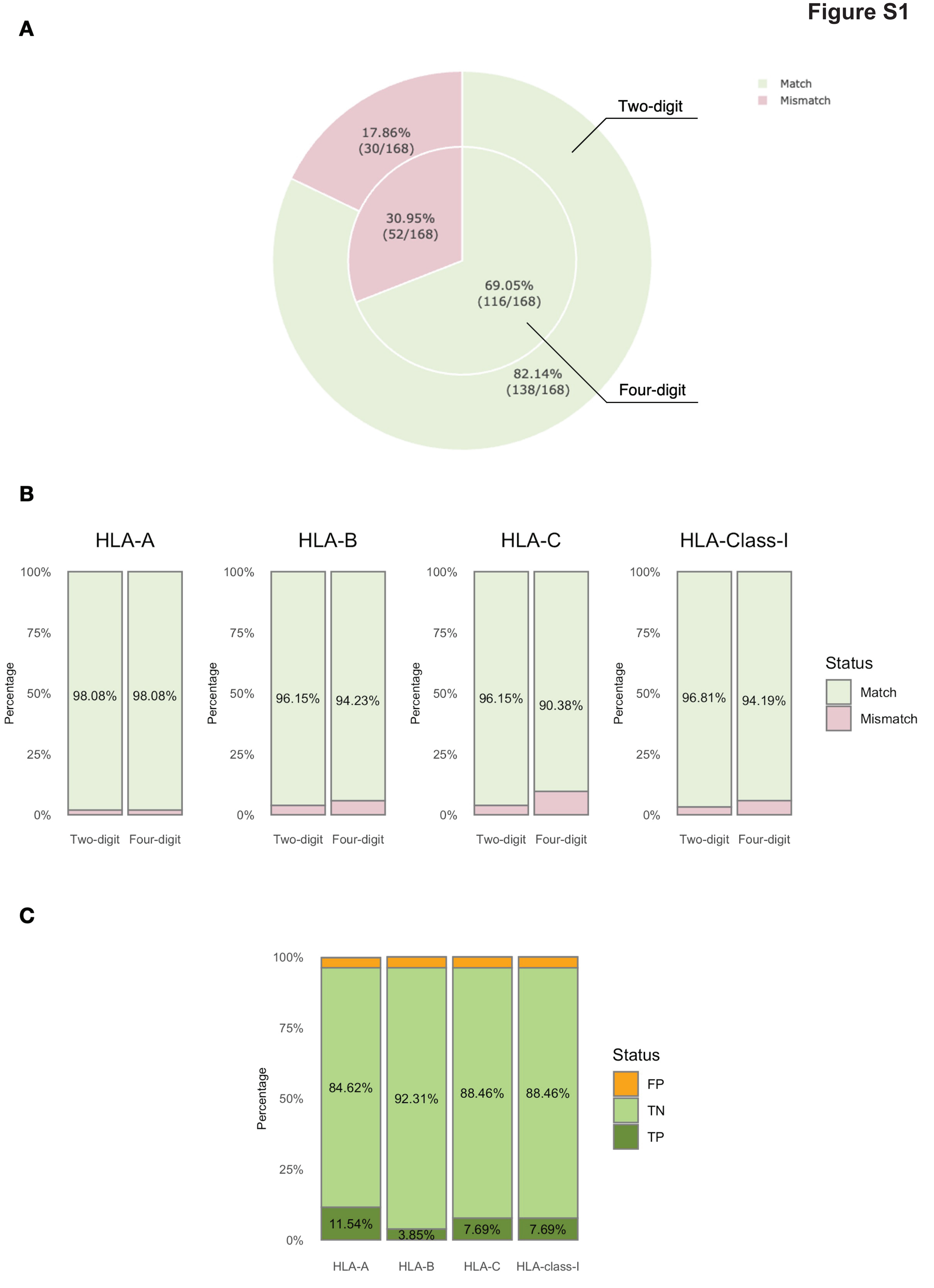

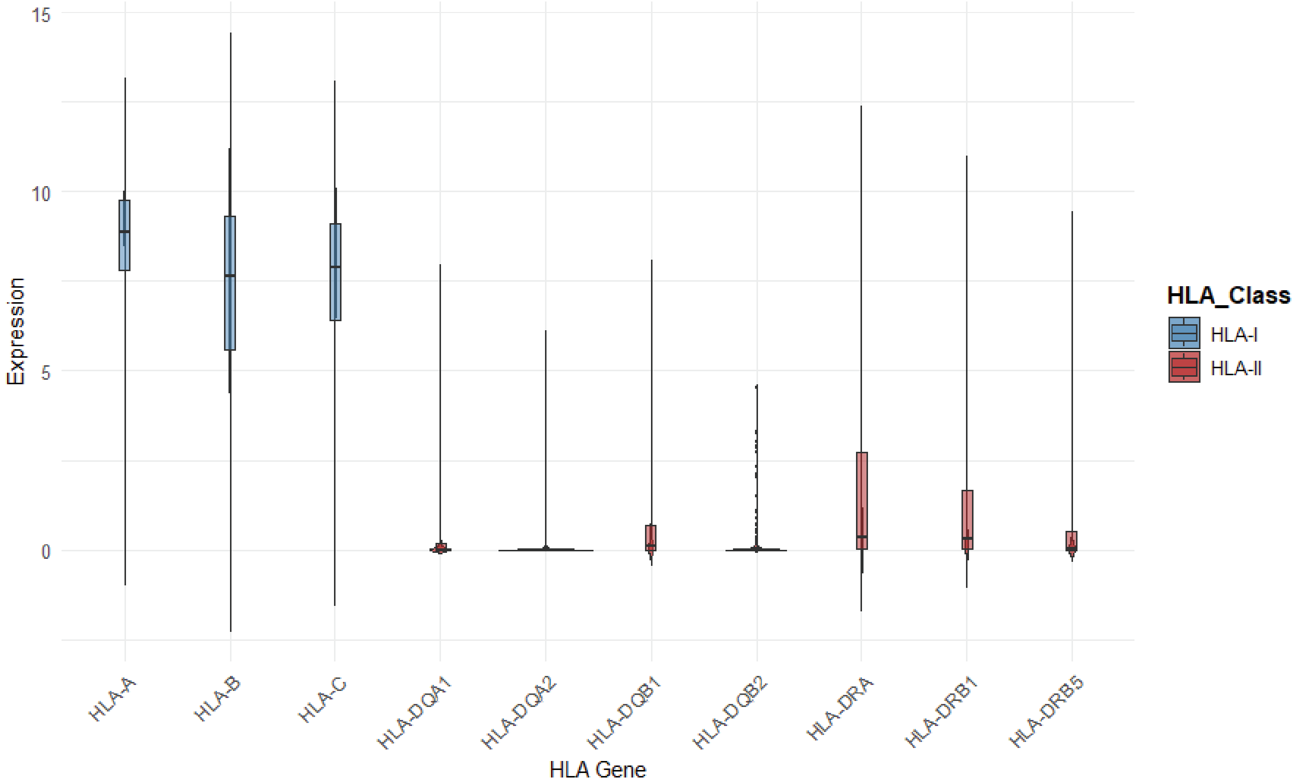

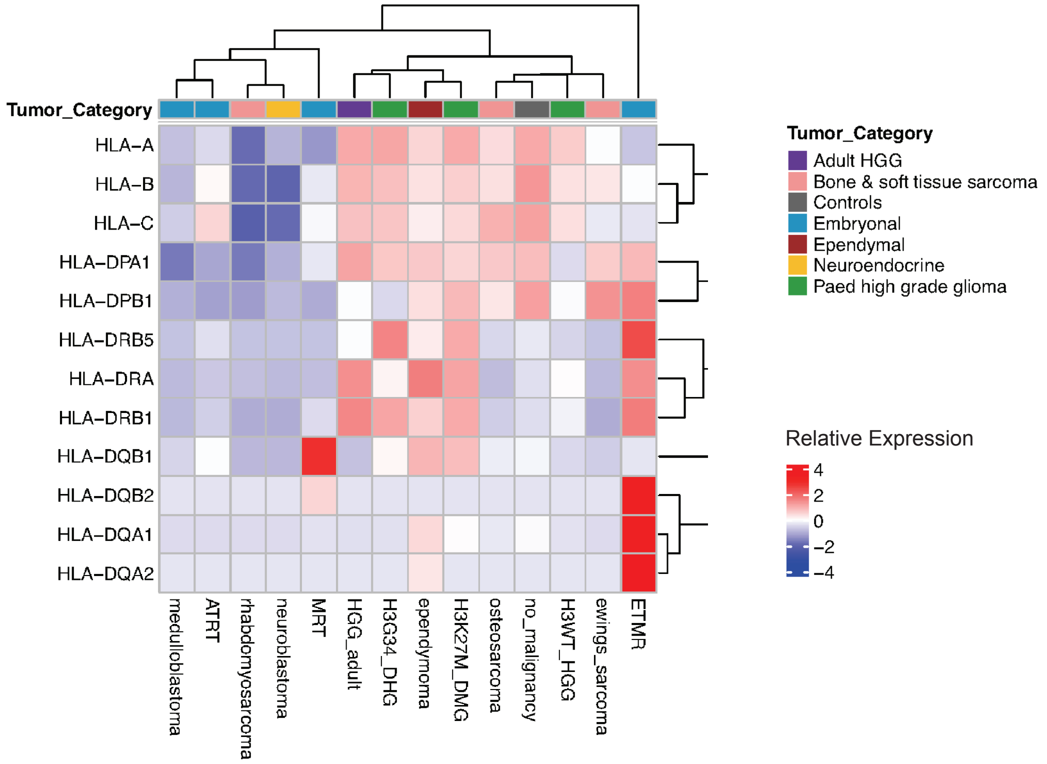

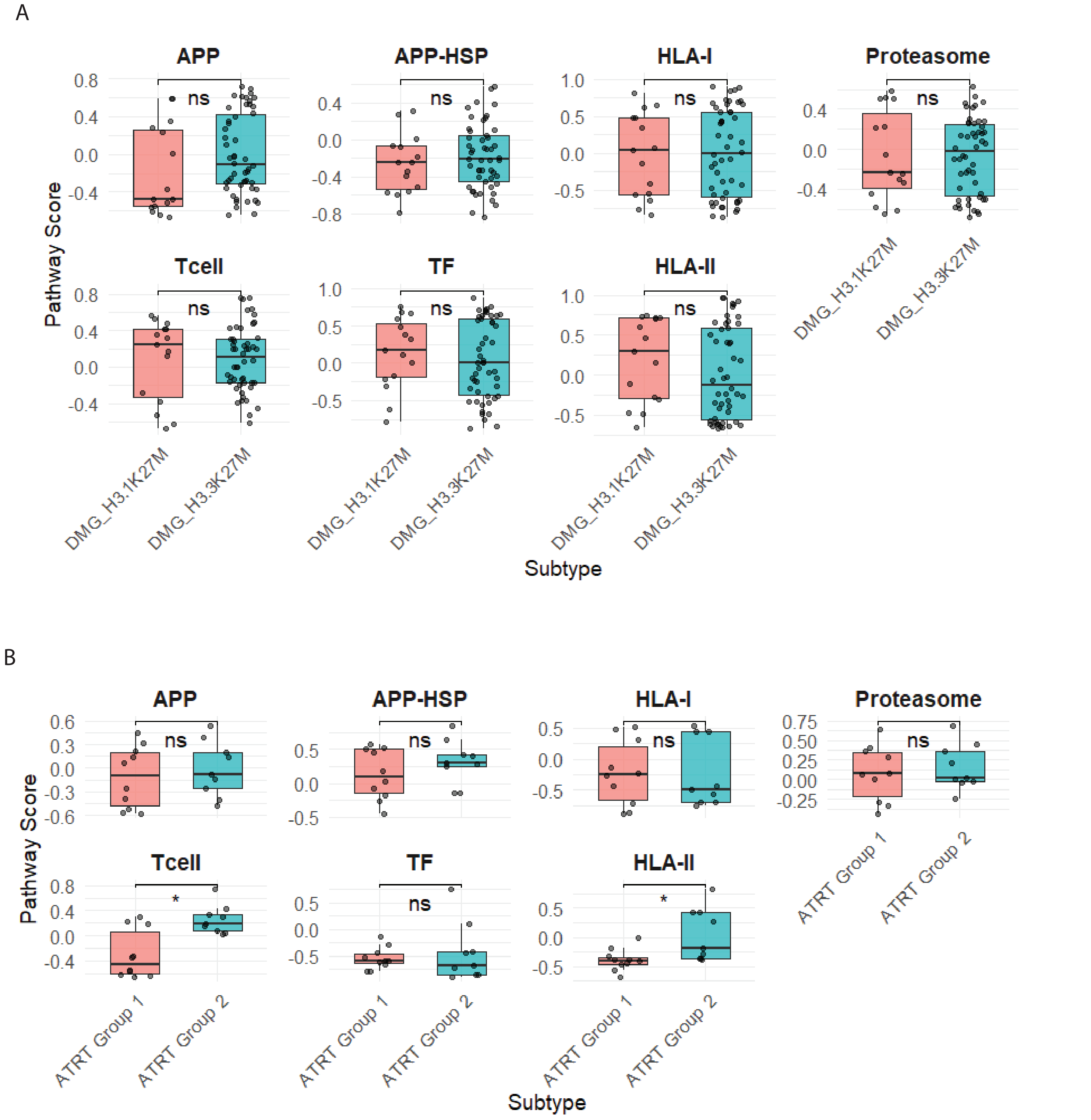

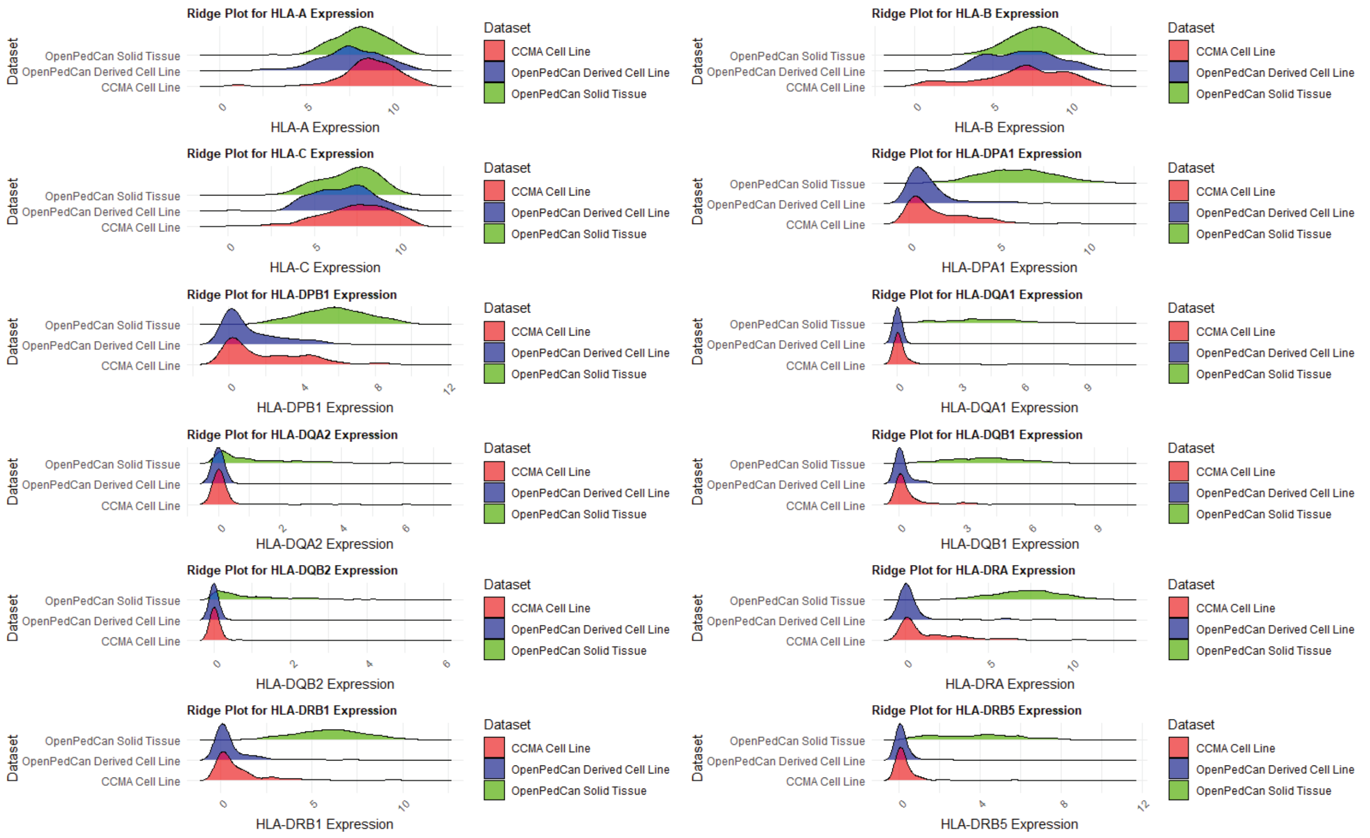

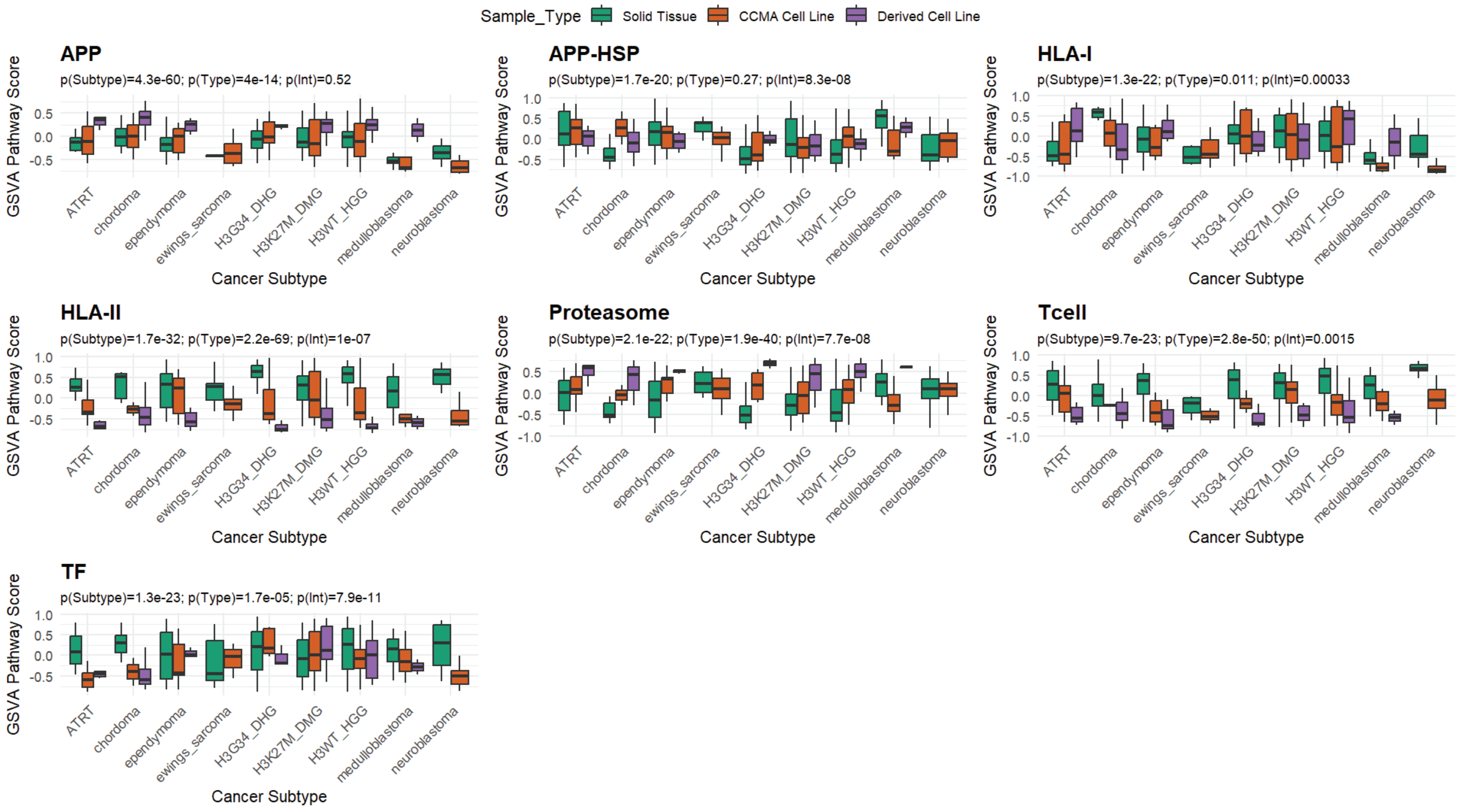
